# Rapid phylogenomic analysis for viral surveillance and metagenomic profiling with Omni2Tree

**DOI:** 10.64898/2026.04.29.721707

**Authors:** Sina Majidian, Adrián Chalco, Xinchang Zheng, Richard J. Webby, Andrew S. Bowman, Rebecca L. Poulson, Nicole M. Nemeth, Fritz J. Sedlazeck, Daniel P. Agustinho

**Affiliations:** Department of Computer Science, Johns Hopkins University, 3400 North Charles St., Baltimore, MD 21218, United States; Human Genome Sequencing Center, Baylor College of Medicine, Houston, TX, USA; Faculty of Science and Engineering, Universidad Peruana Cayetano Heredia, Av. Honorio Delgado 430, Urb Ingeniería, Lima, Peru; St. Jude Children’s Research Hospital, Memphis, TN 38105, USA; The Ohio State University, Columbus, OH 43210, USA; Southeastern Cooperative Wildlife Disease Study, Department of Population Health, College of Veterinary Medicine, University of Georgia, Athens, GA 30602, USA; Department of Pathology, College of Veterinary Medicine, University of Georgia, Athens, GA 30602, USA; Department of Molecular and Human Genetics, Baylor College of Medicine, Houston, TX, USA; Department of Computer Science, Rice University, 6100 Main Street, Houston, TX, USA

## Abstract

Phylogenomic surveillance is limited not by sequencing throughput, but by the difficulty of converting heterogeneous raw data into reliable evolutionary inference, particularly for low-titer and contaminated viral field samples. Here we present Omni2Tree, an assembly-free framework that reconstructs viral phylogenies directly from raw sequencing reads and generates easily shareable interactive reports and genome-wide entropy profiles to identify diversification. In H5N1 benchmark analyses, Omni2Tree maintained accurate placement and topological stability even under low coverage, unlike assembly or reference based methods. Omni2Tree generated an annotated phylogeny for 64-sample H5N1 field surveillance dataset from the eastern USA in under 3 hours. Omni2Tree recovered known phylogenetic structure and key variability insights across 1,328 hepatitis C virus and 707 human cytomegalovirus datasets, and resolved co-infecting respiratory viruses in clinical metagenomic samples. By enabling direct analysis from raw reads, Omni2Tree supports faster, more portable, and more decentralized phylogenomic surveillance across outbreak, clinical, and resource-limited settings.

## Introduction

Phylogenetic trees provide a foundational framework for understanding viral evolution, tracking outbreak dynamics, and identifying therapeutic targets^1^. This has been extensively demonstrated over recent viral outbreaks, such as SARS-CoV-2^2,3^ and Influenza A H5N1^4,5^, which showed that sequencing technologies can generate genomic data at the scale and speed demanded by active outbreaks^6^. However, the computational pipelines required to translate these data into phylogenetic insight have not kept pace, representing a critical and underappreciated bottleneck in genomic surveillance^7^. During the COVID-19 pandemic, these limitations became obvious^8^. While millions of viral genomes were generated globally, only a subset could be incorporated into outbreak monitoring due to constraints in assembly quality, computational scalability, and data integration^9^. Platforms such as Nextstrain^10^ mitigated this challenge through data aggregation and downsampling, enabling centralized global surveillance at the cost of phylogenetic resolution and sample integration, as genomes failing quality thresholds were systematically excluded, leaving a substantial fraction of sequencing data unused. Furthermore, periodic updates for the integration of new samples delay data release as these demand laborious curation, limiting the timeliness of outbreak response.

Current approaches for viral phylogenetics fall into three main categories, each with inherent limitations that are further amplified as typical samples carry low viral titers, host contamination, or population diversity. Assembly-based workflows rely on multi-step pipelines from quality control to tree inference that often necessitate expert curation and centralized infrastructure^11^, and commonly fail when viral burden is low, or host contamination is high, restricting the scope of subsequent phylogenetic studies^12–14^. For segmented viruses such as influenza, de novo assembly frequently produces fragmented assemblies rather than a single contig per segment^15,16^, and concatenation or scaffolding steps introduced to compensate can generate arbitrary chimeric sequences that adversely affect phylogenetic signal^17^. Reference-guided consensus calling addresses some assembly problems, but reconstructed sequences inherit systematic bias from the choice of reference genome, distorting inter-sample relationships inadvertently^18,19^. At the other extreme, alignment-free methods such as Mash^20,21^ offer speed through k-mer sketching but sacrifice the resolution needed to distinguish closely related strains, and in samples with high host contamination, k-mer sketches are dominated by non-target sequence so that distances reflect contamination composition rather than viral phylogeny^22^. Together, these approaches reflect a tradeoff between computational scalability, phylogenetic resolution, and robustness to real-world sample complexity that no existing method resolves.

These limitations are further amplified in multi-pathogen infections, such as respiratory co-infections that impact disease severity and are often undiagnosed^23^. These are frequently occurring in approximately 10-30% of respiratory tract infections in children and a significant portion of adult cases, particularly during peak seasons^24^. Here, the sample itself is effectively a metagenome, where reads originate from distinct organisms simultaneously, and no single reference captures the community^25^. Alternative approaches include de novo genome assembly and binning (e.g., metaSPAdes^26^ and MetaBat^27^) and metagenomic classification methods (e.g., Kraken2^28^), which aim to identify all organisms present in a sample. However, assembly and binning approaches are computationally demanding and sensitive to sequencing depth, require expert curation that is incompatible with outbreak timescales, while classification typically suffers from high false-positive rates^29^, lacks phylogenetic context, and is difficult to reproduce^30^. Approaches explicitly designed for phylogenetic analysis fare no better in this context. They either require pre-built reference trees for placement^31^ or depend on curated marker gene databases^32^, which are not effective for viruses since they lack marker genes.

To overcome these limitations, we present Omni2Tree, a modular phylogenomic framework that reconstructs evolutionary relationships directly from raw sequencing data without assembly or reliance on a single reference genome. Omni2Tree addresses each of the three failure modes described above. It bypasses the assembly step entirely, avoiding fragmentation and chimeric artifacts. It constructs orthologous groups de novo from user-provided reference assemblies, identifying genes shared by at least two members of the database, and maps reads against these rather than against a single reference, so that tree topology is not anchored to any one genome. And it operates natively on metagenomic samples containing multiple co-infecting pathogens in a single analysis. Omni2Tree places raw reads from short- and long-read technologies alongside genome assemblies on a unified phylogeny, eliminating the need to choose between input types. While our previous work, Read2Tree ^7^, enabled assembly-free phylogenetics for single-organism data, Omni2Tree extends this to viral and metagenomic applications at scale. Beyond tree inference, the same run produces interactive metadata-aware visualization and per-position entropy profiles across the genome, providing both phylogenetic context and diversity information without additional analysis steps. We benchmark Omni2Tree against assembly-based, reference-guided, and alignment-free approaches across nine coverage levels on H5N1 outbreak samples, demonstrate scalability on over 2,000 HCV and hCMV samples, and validate metagenomic strain-level resolution on viral co-infection samples and the CAMI benchmark.

## Results

### Omni2Tree enables assembly-free phylogenomics across viral and metagenomic applications

Omni2Tree is an assembly-free phylogenomic framework that uses raw sequencing reads to enable direct inference of maximum-likelihood phylogenetic trees with integrated metadata visualization and evolutionary diversity analysis (**Figure 1**). Omni2Tree is read-length and sequencing technology agnostic, allowing the user to process both short- and long-read samples, as well as assembled genomes simultaneously. Omni2Tree is compatible with both RNA and DNA viruses, demonstrated here across influenza A (segmented RNA), Hepatitis C virus (positive-sense RNA), human Cytomegalovirus (dsDNA), and mixed viral metagenomes containing co-infections. There are three main steps in Omni2Tree: i) Automated marker gene construction and alignment based on user-provided assemblies (**Figure 1A**). For segmented viruses, the analysis can be restricted to individual segments or subsets of segments, allowing users to investigate reassortment or focus on genes of particular interest. ii) Processing of the individual samples by aligning to the generated marker genes and sequence reconstruction (**Figure 1B**). Because each sample is processed independently, this can scale to thousands of samples analyzed in parallel and enables the integration of new samples at a later stage. iii) Tree inference and visualization (**Figure 1C**). By default, Omni2Tree performs bootstrapped maximum-likelihood inference using IQ-TREE2, but it also produces a concatenated alignment of all reconstructed genes from samples and references. This alignment can serve as input for alternative inference methods, including Bayesian approaches such as BEAST2^33^, giving users flexibility in their choice of phylogenetic framework. Omni2Tree software is publicly available as an open-source package at https://github.com/DanielPAagustinho/omni2tree.

**Figure 1.**
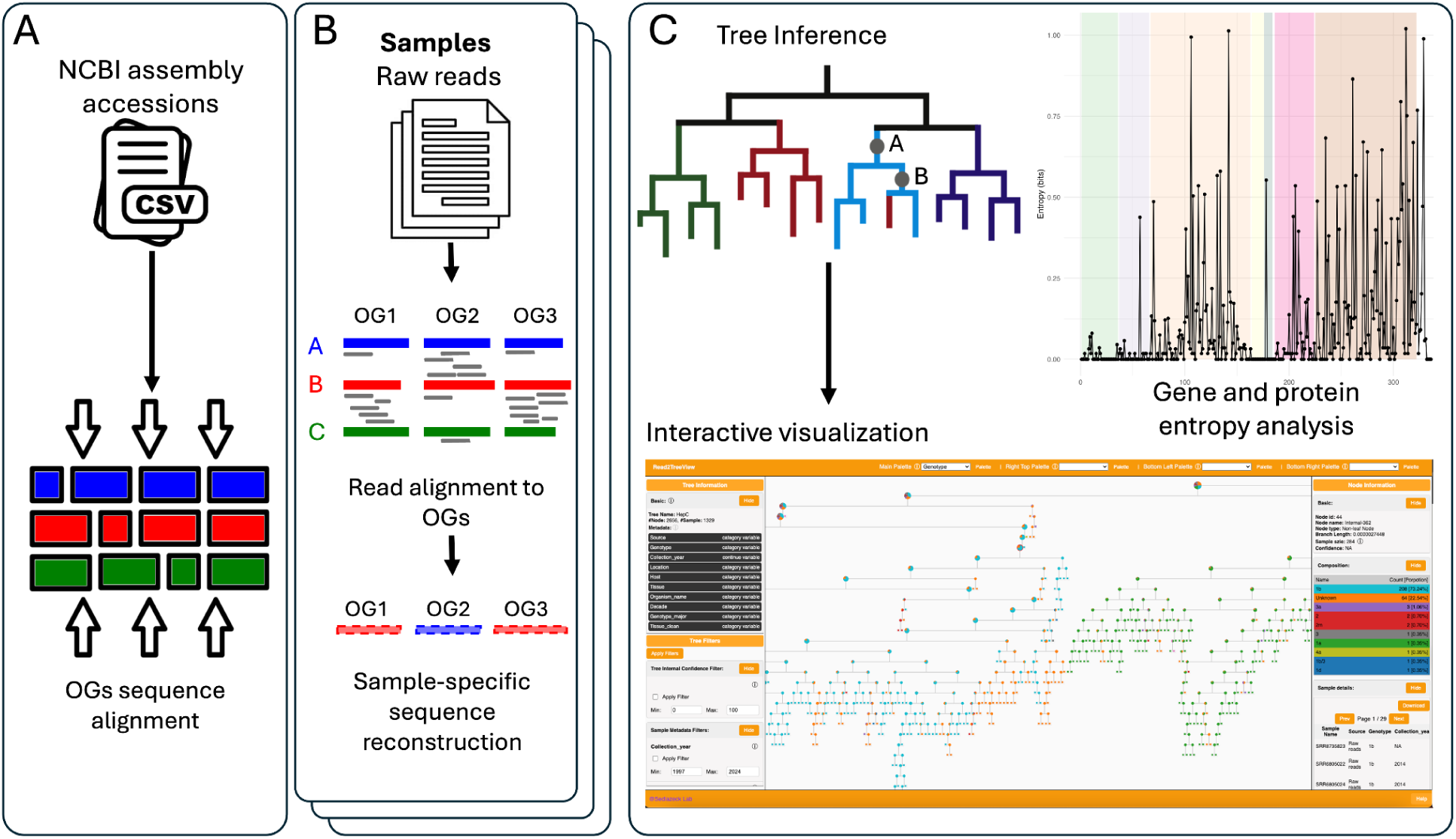
Omni2Tree workflow overview. **(A)** A list of NCBI accessions is used to download coding sequences and construct a marker gene reference database via orthologous group (OG) inference, producing per-gene multiple sequence alignments shared across all subsequent analyses. **(B)** Raw sequencing reads from each sample are downloaded and mapped independently to the OG references, and a sample-specific representative sequence is reconstructed per OG from the mapped reads. Optional deduplication and downsampling are applied before mapping. Omni2Tree operates in two modes: in isolate mode, shown here, the sequence most supported by reads is chosen per OG, while in metagenomic mode, multiple sequences are reconstructed per OG to capture reads originating from different species. Because samples are processed independently, Step 2 scales to thousands of samples in parallel, and new samples can be added to an existing analysis without reprocessing prior data. **(C)** A metadata table is combined with the Step 1-2 outputs to produce three integrated outputs. A bootstrapped maximum-likelihood tree is inferred from the concatenated multi-OG alignment (IQ-TREE2). The tree is rendered in Omni2TreeView, a self-contained interactive HTML small file (≤1 MB for hundreds of samples), readable offline without additional software. It supports filtering and coloring by any metadata attribute available, with per-node and per-sample details accessible on demand. Per-gene and per-protein Shannon entropy profiles are calculated across all samples, reported as publication-ready plots with optional functional domain annotation, identifying positions under immune selection pressure or functional constraint. Reference assemblies (Step 1) and read-derived sequences (Step 2) are placed on the same tree, enabling unified analysis of assembled and unassembled data in a single run.

Omni2Tree outputs an interactive HTML file (Omni2TreeView) that renders the inferred tree alongside reference sequences and allows dynamic filtering and coloring by any selection of metadata classes (genotype, geography, collection date, tissue, or any user-defined attribute) in a single file. The resulting file is self-contained, requires no server or additional software to open, and remains under 1 MB for datasets of several hundred samples, making it shareable in low-bandwidth or resource-limited surveillance settings. Shannon entropy is calculated per-position across all processed samples, and reported as publication-ready per-gene plots for both nucleotide and amino acid sequences, providing an immediate summary of diversity across the genome without additional analysis steps. Together, these outputs take a dataset from raw reads to annotated phylogeny, diversity profiles, and interactive visualization in a single workflow requiring no manual curation between steps.

### Omni2Tree maintains phylogenetic accuracy in the presence of contamination

A panel of 21 H5N1 short-read samples alongside 11 reference assemblies served as the primary benchmark for Omni2Tree against assembly-based and reference-guided consensus approaches. (**Supplementary Tables 1 and 2**). For the purposes of this benchmark, we are analyzing whole genomes instead of any specific segment. Twenty of the raw-read samples were collected from different hosts during the 2024 US dairy cattle outbreak (clade 2.3.4.4b; BioProject PRJNA1102327)^4^, and sample SRR11611114 (clade 2.3.2.1c) serves as a phylogenetic placement control present as both raw reads and curated genome assembly, representing typical low-viral-burden field samples^34^ with only 0.31% H5N1 reads. Viral read fractions for all 21 samples are reported in **Supplementary Table 1**. Systematic downsampling from full sequencing depth (median coverage of 3,155x) to 0.5x coverage was used to assess how each approach degrades under data-limited conditions. We compared Omni2Tree with three other widely used approaches: (1) de novo assembly with MegaHit; (2) reference-guided alignment and consensus calling with BWA-MEM^35^ and bcftools^36^, using two reference genomes, a closely related cattle isolate (2024) and a divergent avian isolate (2022) from the same 2.3.4.4b cluster (**Supplementary Table 2**); (3) alignment-free k-mer phylogenetics with MashTree.

Patristic distance (i.e., sum of the branch lengths connecting two nodes in a phylogenetic tree) to the cognate assembly served as the primary measure of placement accuracy for SRR11611114 across approaches and coverage levels. Omni2Tree correctly placed the sample, achieving near-zero patristic distance (∼0.000002) to its cognate assembly (**Figure 2A**) at full coverage. MegaHit + IQ-TREE placed it furthest from the cognate assembly (patristic distance ∼0.95), as expected from the assembly of predominantly contaminant reads. Assembly quality degraded substantially and non-linearly with decreasing coverage. At full sequencing depth, MegaHit produced a total assembly size with inflated contig numbers (**Supplementary Figure 1A**), and frequently exceeding the 13.6 kb reference genome length, often by more than 200% (**Supplementary Figure 1B**). This over-assembly likely reflects DNA contamination that MegaHit cannot distinguish from viral sequence. As expected, the assemblies became increasingly fragmented and incomplete at 2x coverage and below, with median sizes falling below 50% of the expected genome length (**Supplementary Figure 1B**). MashTree recovered the correct topological placement of SRR11611114 alongside its cognate assembly, but with a patristic distance of ∼0.25 rather than near zero (**Figure 2A**). Although MashTree branch lengths reflect MinHash sketch distances rather than substitutions per site and are not directly comparable to the other approaches in absolute terms, the expected distance between a sample and its own cognate assembly is still zero under any distance metric. The observed displacement indicates that host contamination diluted the viral k-mer signal enough to erode the similarity between sample and reference, even when topological placement remained correct. Reference-guided consensus calling avoided assembly fragmentation but introduced reference genome bias as a systematic failure. When the cattle Texas reference was used, the patristic distance from SRR11611114 to its cognate flamingo assembly appeared low (∼0.015 vs ∼0.829 with the Eagle reference), superficially suggesting correct placement (**Figure 2A**). This apparent proximity reflects reference-induced compression rather than true phylogenetic signal, as consensus sequences generated by mapping to the cattle reference are anchored to it at low-coverage positions, artificially collapsing inter-sample distance.

**Figure 2.**
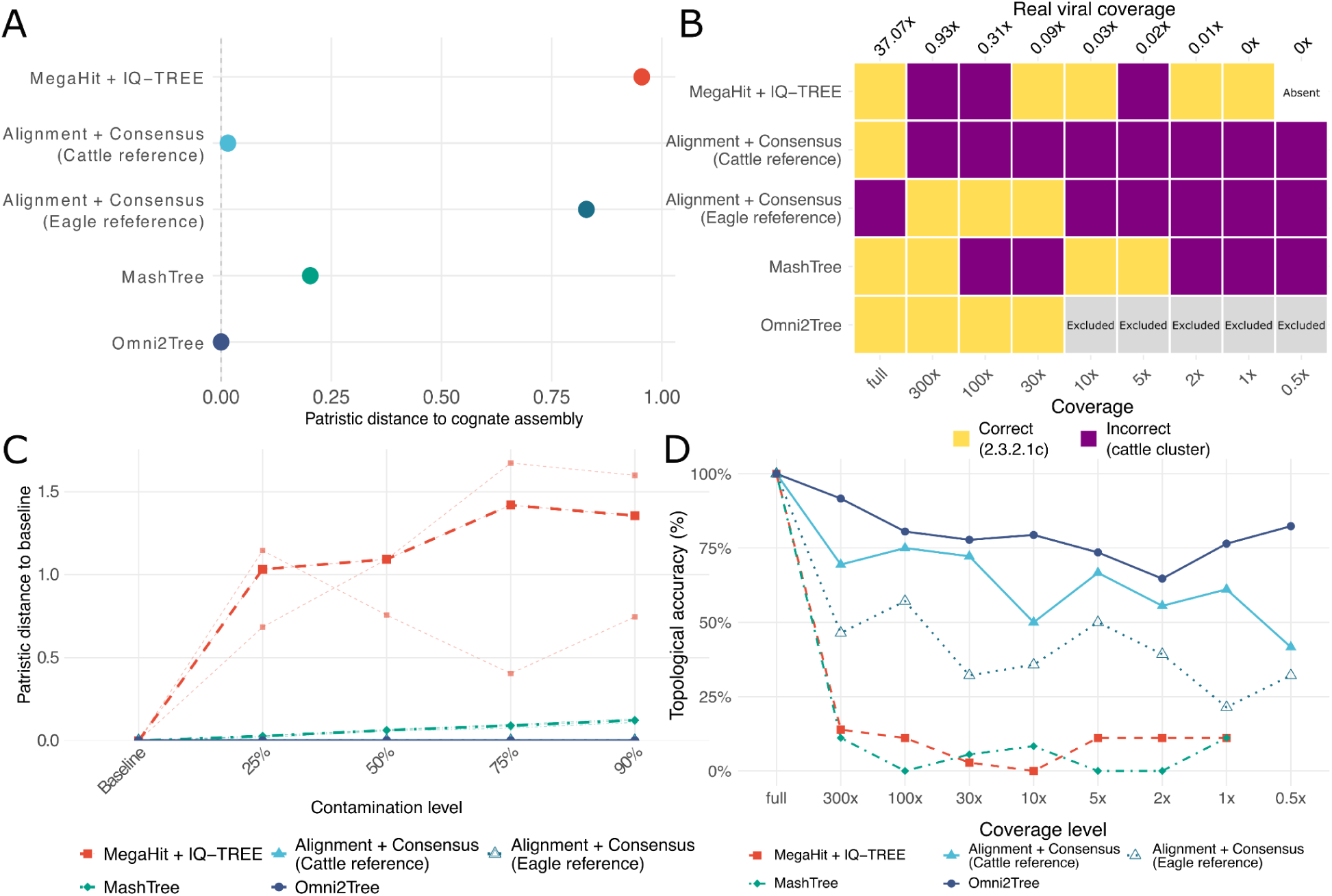
Omni2Tree maintains phylogenetic accuracy and topological stability across coverage levels. **(A)** Patristic distance from SRR11611114 (clade 2.3.2.1c raw reads, full coverage) to its cognate assembly (A/flamingo/Kazakhstan/6570/2015) in trees produced by each approach. Zero indicates perfect co-placement with the cognate assembly. **(B)** Clade membership of SRR11611114 across approaches and coverage levels. Correct placement indicates SRR11611114 was closer to its cognate assembly (A/flamingo/Kazakhstan/6570/2015) than to the 20 cattle outbreak samples. Coverage labels show nominal subsampling depth and estimated real Influenza A coverage for SRR11611114 (0.31% of reads). White: sample absent from the tree. Grey: Omni2Tree actively excluded samples with insufficient viral reads after downsampling. **(C) Effect of host contamination on phylogenetic placement across approaches.** Patristic distance from each contaminated version of a sample to its uncontaminated baseline, measured within a single combined phylogenetic tree per approach. Bovine genomic reads were spiked into three high-purity H5N1 samples at contamination levels of 25%, 50%, 75%, and 90% of total reads. Thin lines show individual samples; thick lines show the median across the three samples. A distance of zero indicates that contamination did not alter phylogenetic placement. Distances for the assembly, alignment + consensus and Omni2Tree are measured in substitutions/site, while Mashtree branch lengths represent Mash distances derived from k-mer similarity and are not directly comparable in absolute magnitude to substitution-based distances. However, within each approach, the relative change from baseline to contaminated conditions remains interpretable.**(D)** Within-approach topological stability: Robinson-Foulds distance of each subsampled tree against the same approach’s full-coverage tree, restricted to outbreak samples only (n=20). Higher values indicate greater topological consistency with the full-coverage result. Bootstrap support ≤70 collapsed prior to comparison.

The topological placement of SRR11611114 within the 2.3.2.1c or with the cattle (2.3.4.4b) cluster varies depending on the sequencing coverage values (**Supplementary Figures 2**-**6)**. Omni2Tree was able to place SRR11611114 correctly alongside its cognate assembly, with bootstrap support of 100 and a long separating branch from the 2.3.4.4b cluster (**Figure 2A**, **Supplementary Figure 6)**. But as coverage decreases, real viral coverage becomes very low. Omni2Tree was able to place SRR11611114 correctly in a cluster with its own cognate assembly down to 30x coverage, when we expect only 0.093x coverage of true H5N1 reads across this sample due to the high contamination. At lower coverages, Omni2Tree correctly excluded this sample from being placed into the phylogeny (**Figure 2B**). This is in contrast to the other approaches, where assembly and consensus-based methods both attempted to place the sample despite an insufficient viral signal, consistently producing incorrect placements. The alignment + consensus approach using the cattle reference placed SRR11611114 within the incorrect cattle (2.3.4.4b) cluster at 300x or lower coverages. In contrast, MegaHit, MashTree, and the eagle reference approach placed SRR11611114 randomly in 2.3.4.4b and 2.3.2.1c clusters as coverage varied, indicating noise-driven rather than systematically biased failures. Thus, in contrast to Omni2Tree, the other approaches produced topologically incorrect placements without any indication of data insufficiency (**Figure 2A**, **Supplementary Figure 2-7**).

To quantify robustness to host contamination, we artificially generated contaminated samples. To do so, we combined increasing amounts of bovine genomic reads into the three highest purity H5N1 cattle outbreak samples, with final contamination levels ranging from 25% to 90% of total reads. We measured the patristic distance between baseline and different levels of contaminated versions of each sample in a combined phylogenetic tree (**Figure 2C**). Reference-guided consensus calling and Omni2Tree were unaffected by contamination across all levels tested. Both approaches successfully filter or ignore non-viral sequences by design, either through reference-guided mapping or marker gene alignment. MashTree showed increasing degradation proportional to contamination level, as k-mer sketches incorporate host sequence without discrimination. As expected, MegaHit assembly showed the largest patristic distances with high variability between the samples (**Figure 2C**, dashed lines), consistent with the assembly of highly contaminated samples without extra purification steps. This clearly shows the failure of the assembly approach in stark contrast to the successful performance of reference-guided calling and Omni2Tree.

### Omni2Tree maintains phylogenetic stability at decreasing coverage levels

Having established placement accuracy for a single known sample (SRR11611114), we next asked whether the inferred topology across the 20 cattle outbreak samples remains consistent as coverage degrades, using within-approach Robinson-Foulds distances against each method’s own full-coverage tree. Omni2Tree maintained the highest topological stability across all coverage levels, remaining above 75% identity with its own full-coverage tree at most conditions tested, including 0.5x (**Figure 2D**). In contrast, MegaHit + IQ-TREE collapsed to <20% topological identity already by 300x coverage, and MashTree showed similarly rapid degradation. The progressive loss of assembly completeness (**Supplementary Figure 1**) directly explains MegaHit’s topological collapse. The alignment + consensus approach with the cattle reference maintained intermediate stability (∼40-65% across subsampled conditions), but this stability is partly artificial, as shown above, consensus sequences are anchored to the reference regardless of coverage, so the tree topology is constrained by the reference rather than by the data. At low coverage, this approach calls fewer variants and fills more positions with the reference sequence, causing consensus sequences to converge toward the reference chosen regardless of the sample’s true biology. This reference dependence is not limited to low-coverage conditions. Even at full sequencing depth, the choice of reference genome fundamentally altered the topology of the outbreak samples. Consensus trees built with the alignments to the cattle and eagle references produced different sample clustering despite identical input reads (**Supplementary Figure 7**), confirming that the recovered phylogeny reflects the reference as much as the underlying biology.

Omni2Tree’s computational requirements were comparable to reference-guided consensus calling. At full coverage, Omni2Tree required 55.8 CPU hours (51.8 hours without step 1), versus 12.6 CPU hours for the alignment + consensus pipelines. MegaHit + IQ-TREE required 653.0 CPU hours. MashTree was the fastest approach at 1.5 CPU hours.

### Omni2Tree extends to long-read sequencing and detects samples with insufficient viral coverage

To showcase Omin2Tree also works independent of the sequencing technology we analyzed 22 RSV/A samples from PRJNA980575 sequenced with Oxford Nanopore Technology (ONT, **Supplementary Table 4**). These samples have an average read length of 300-700 bp, which is expected from amplicon-based sequencing protocols commonly used in respiratory surveillance^37^. Omni2Tree processed the same raw reads directly, placing 21 of the 22 samples in a phylogenetic tree. Upon inspection, the excluded sample, SRR24833889, showed zero reads originating from RSV, as determined by read alignment to an RSV reference genome (**Supplementary Table 3**). All 21 valid samples were placed within the RSV/A clade and outside the RSV/B clade, confirming that subgroup assignment is driven by read content rather than by the composition of the reference panel.

Standard long-read assembly tools such as Flye^38^ and Canu^39^ failed on these datasets due to insufficient read length. To generate reference phylogenies, we constructed reference-guided consensus sequences by aligning with minimap2 to an RSV/A reference genome, followed by variant calling^36^ and consensus generation (see methods). This approach placed all 22 samples in the tree regardless of per-sample read depth. Thus, sample SRR24833889 was wrongly placed by this method despite the complete absence of RSV reads, producing a consensus sequence, and thus an erroneous phylogenetic placement.

Phylogenetic trees generated by Omni2Tree and the alignment + consensus approaches are shown in **Supplementary Figure 8**. Together, these results show that Omni2Tree processes ONT data without parameter tuning, and that its quality threshold provides a safeguard against placements unsupported by read evidence, a failure mode that reference-guided consensus approaches do not expose.

### Rapid phylogenetic surveillance of emerging H5N1 outbreak samples

To evaluate Omni2Tree in a real-world surveillance setting, we analyzed 64 H5N1 samples provided by the St. Jude Center of Excellence for Influenza Research and Response, collected through surveillance and outbreak response efforts across the eastern United States. Bovine samples were obtained from an infected Ohio dairy farm early in the 2024 outbreak, while wildlife samples were collected primarily through passive surveillance of dead or moribund animals submitted to the Southeastern Cooperative Wildlife Disease Study, University of Georgia (**Figure 6, Supplementary Table 5**). The dataset spans 17 host species, including 18 bovine samples and 46 wildlife samples from 8 states (Georgia, Kansas, Kentucky, Nebraska, North Carolina, Ohio, South Carolina, and Virginia), collected between 2023 and 2024. Forty samples originated from avian hosts (ducks, geese, raptors, grebes, and other wild birds), while 24 came from mammalian hosts (cattle, raccoons, a black bear, and a mountain lion). Several animals were sampled from multiple tissues, including a raccoon (nasal, oral, and rectal swabs) and a redhead duck (brain, oropharyngeal, adrenal gland, and heart swabs), providing within-host sampling depth. Four reference assemblies spanning 2002 to 2024 were included as phylogenetic context. For several isolates, both oropharyngeal swabs (OP, collected directly from the animal) and subsequent egg isolates (E1) generated from these swabs were sequenced independently, providing biological replicates that allow assessment of mutations accumulated during laboratory passage and representing different sequence qualities.

Omni2Tree processed all 64 samples from raw reads to a publication-ready annotated phylogeny in under 3 hours using 5 CPU cores and 16 GB RAM. The resulting tree (**Figure 3**) separated bovine and wildlife samples into distinct clusters consistent with their epidemiological context and suspected genotypic diversity. The Ohio dairy cattle samples formed a tight clade alongside the 2024 commercial milk reference, as expected for an ongoing outbreak with limited intra-herd diversity. Wildlife samples showed considerably more phylogenetic diversity, spanning collections from 2023 and 2024 across multiple states. The greater phylogenetic diversity among wildlife samples likely reflects frequent reassortment events, which are common in wild bird populations where co-circulation of multiple viral lineages and subtypes provides opportunities for segment exchange. Mammalian spillover samples (raccoon, black bear, mountain lion) occupied longer terminal branches, consistent with both independent cross-species transmission from a diverse avian pool and reduced reassortment once in mammalian hosts. The tight clustering of the Ohio dairy cattle samples, by contrast, is consistent with limited reassortment during sustained cattle-to-cattle transmission within the same geographical and collection date constraints. Multi-tissue samples from the same animal clustered together, and OP/E1 pairs from the same isolate showed minimal divergence, confirming that Omni2Tree produces consistent phylogenetic placement regardless of sample type or passage history.

**Figure 3:**
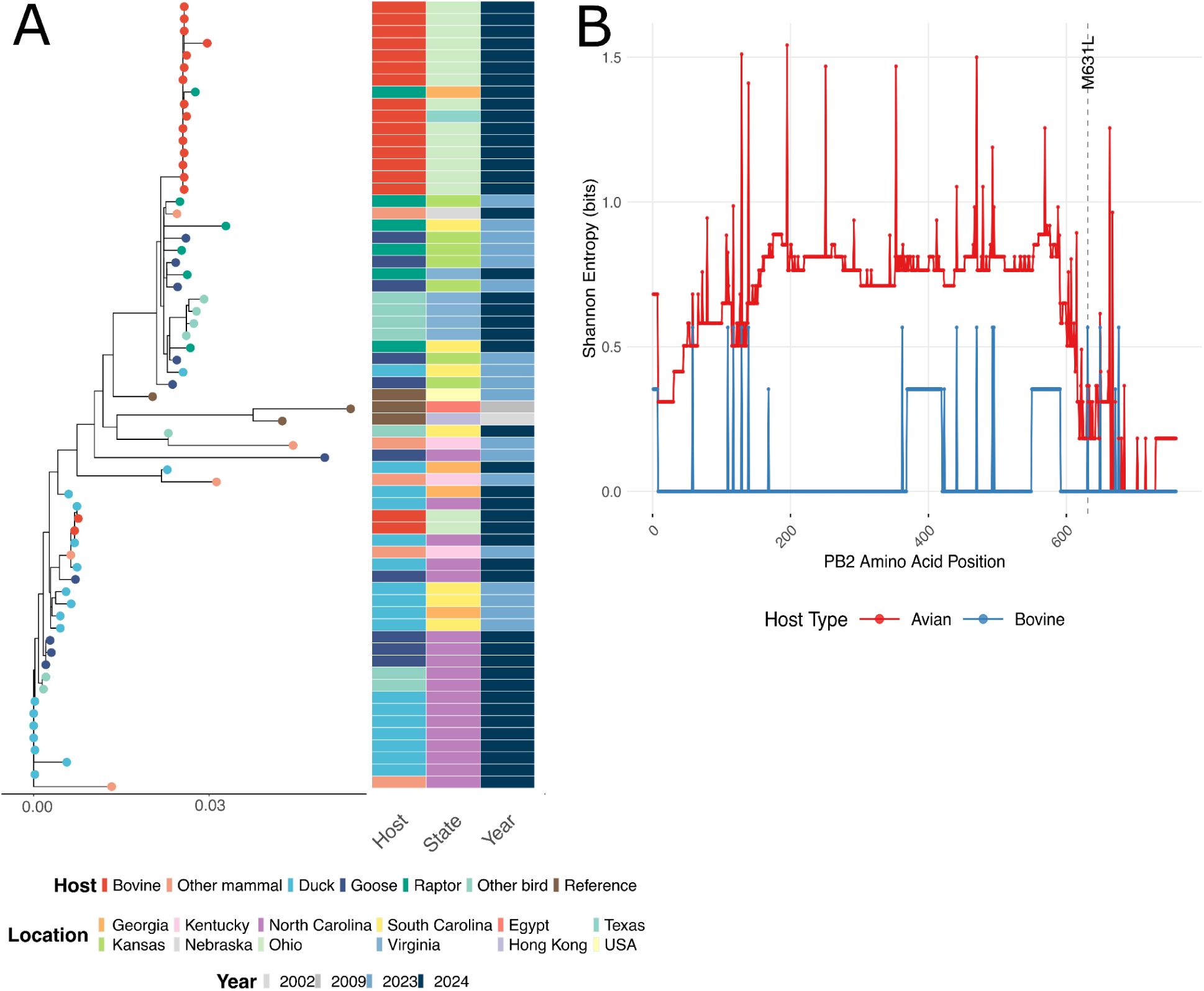
Omni2Tree phylogenomic analysis of 64 H5N1 samples from an active surveillance effort across the eastern United States. **(A)** Maximum-likelihood phylogeny inferred directly from raw sequencing reads using Omni2Tree, with four reference assemblies included as phylogenetic context (2002-2024). Tip points are colored by host groups. The three annotation strips indicate, from left to right, host group, US state of collection, and year of collection. The analysis, from raw FASTQ files to annotated phylogeny, completed in under 3 hours using 5 CPU cores and 16 GB RAM. The scale bar represents substitutions per site. **(B)** Shannon entropy (bits) was calculated at each reference-mapped amino acid position for avian (red, n=36) and bovine (blue, n=15) sample groups using Omni2Tree’s entropy module. Higher entropy indicates greater sequence diversity at a given position. The bovine samples show near-zero entropy across most of the protein, consistent with clonal spread from a limited introduction event, while avian samples display elevated entropy throughout, reflecting the broader diversity of H5N1 circulating in wild bird populations. The dashed line marks position 631, where an M631L substitution was observed in 13 of 15 bovine samples but only 1 of 36 avian samples. The canonical mammalian adaptation markers E627K and D701N were absent in both groups.

Taking advantage of Omni2Tree’s metadata-based grouping feature, we compared entropy profiles between cattle-derived and avian-derived samples (**Supplementary Figure 9**). Avian samples showed consistently higher entropy across nearly all proteins, reflecting the broader phylogenetic diversity of H5N1 circulating in wild bird populations, where reassortment and co-circulation of multiple lineages maintain substantial sequence variation. Cattle samples, by contrast, displayed markedly lower entropy, with long stretches of near-zero variation particularly evident in the polymerase complex proteins (PB1, PB2, PA) and NP. This is consistent with the cattle outbreak representing a single or very few introductions from the avian reservoir, followed by sustained cattle-to-cattle transmission with limited further reassortment, effectively sampling a narrow slice of the broader clade diversity. The difference was less pronounced for the surface glycoproteins HA and NA and for NS1, where cattle samples retained moderate entropy at several positions, possibly reflecting ongoing immune-driven selection even within the constrained cattle lineage. These entropy profiles complement the phylogenetic analysis (**Figure 3**) and reinforce the interpretation that the Ohio dairy outbreak originated from a limited spillover event, with subsequent clonal spread within the herd.

To examine this further, we focused on PB2 (**Figure 3B**), the protein most directly implicated in mammalian adaptation of avian influenza^40^. The entropy contrast between avian and bovine samples was particularly striking across the length of the protein, with bovine samples showing near-zero entropy over long stretches. We screened for known mammalian adaptation markers, including E627K and D701N, the two canonical substitutions associated with enhanced polymerase activity at mammalian body temperatures^40^. Consistent with previous reports from the 2024 North American outbreak^41^, neither substitution was present in our bovine or avian samples. However, we identified M631L in 13 of 15 bovine samples but only 1 of 36 avian samples. M631L has been reported as a compensatory mutation that can enhance polymerase activity in the absence of E627K^41^, suggesting that this cattle lineage may be achieving mammalian adaptation through an alternative molecular pathway.

This analysis demonstrates that Omni2Tree can deliver actionable phylogenetic results from a heterogeneous field surveillance dataset within hours, a turnaround that would allow near-real-time tracking of viral spread across host species and geographic regions during an active outbreak. The additional entropy profiles complement the phylogenetic analysis (**Figure 3A**) and reinforce the interpretation that the Ohio dairy outbreak originated from a limited spillover event, with subsequent clonal spread and potential host-specific adaptation within the herd.

### Omni2Tree scales to thousands of samples across divergent RNA viral datasets

To demonstrate Omni2Tree’s capacity to handle large, heterogeneous viral datasets, we applied it to 1,328 Hepatitis C virus (HCV) samples comprising 613 genome assemblies and 715 raw read datasets (**Supplementary tables 6** and **7**). HCV is a highly diverse positive-sense RNA virus whose seven genotypes differ by up to 30% at the nucleotide level^42^, providing a stringent test across deep divergences. The resulting phylogeny (**Figure 4A**) showed clear separation of the seven major genotypes into distinct clades, with genotype serving as the strongest predictor of phylogenetic clustering (Cramér’s V = 0.80). Geographic substructure was evident within individual genotypes (GT1-7). GT4 samples were predominantly from the Netherlands, GT6 from Hong Kong and Southeast Asia, and GT3 from the UK and South/Southeast Asia, in accord with known epidemiological distributions. Within GT1, the most geographically widespread genotype, subtype GT1a predominated in North American and German samples while GT1b was more prevalent in Spain and Switzerland, recapitulating established transmission patterns^43,44^. These multi-layered patterns of genotype, geography, and subtype are difficult to represent simultaneously in a conventional tree visualization. **Figure 4B** shows the same tree rendered through Omni2TreeView, which supports concurrent display of multiple metadata classes and makes these associations directly interpretable. Specifically, a subset of 32 readsets showed discordant phylogenetic placement relative to their annotated genotype (white tip symbols, **Figure 4A**). These outliers are consistent with assay-constrained inputs that provide incomplete phylogenomic signals, as commonly encountered in bulk SRA retrieval, including single-locus NS5B resistance amplicons^45–48^ and earlier probe designs^49^. No systematic clustering by data source was observed, assemblies and raw read datasets were distributed throughout all genotypic clades, confirming that Omni2Tree’s consensus sequences are phylogenetically equivalent to genome assemblies regardless of input format.

**Figure 4.**
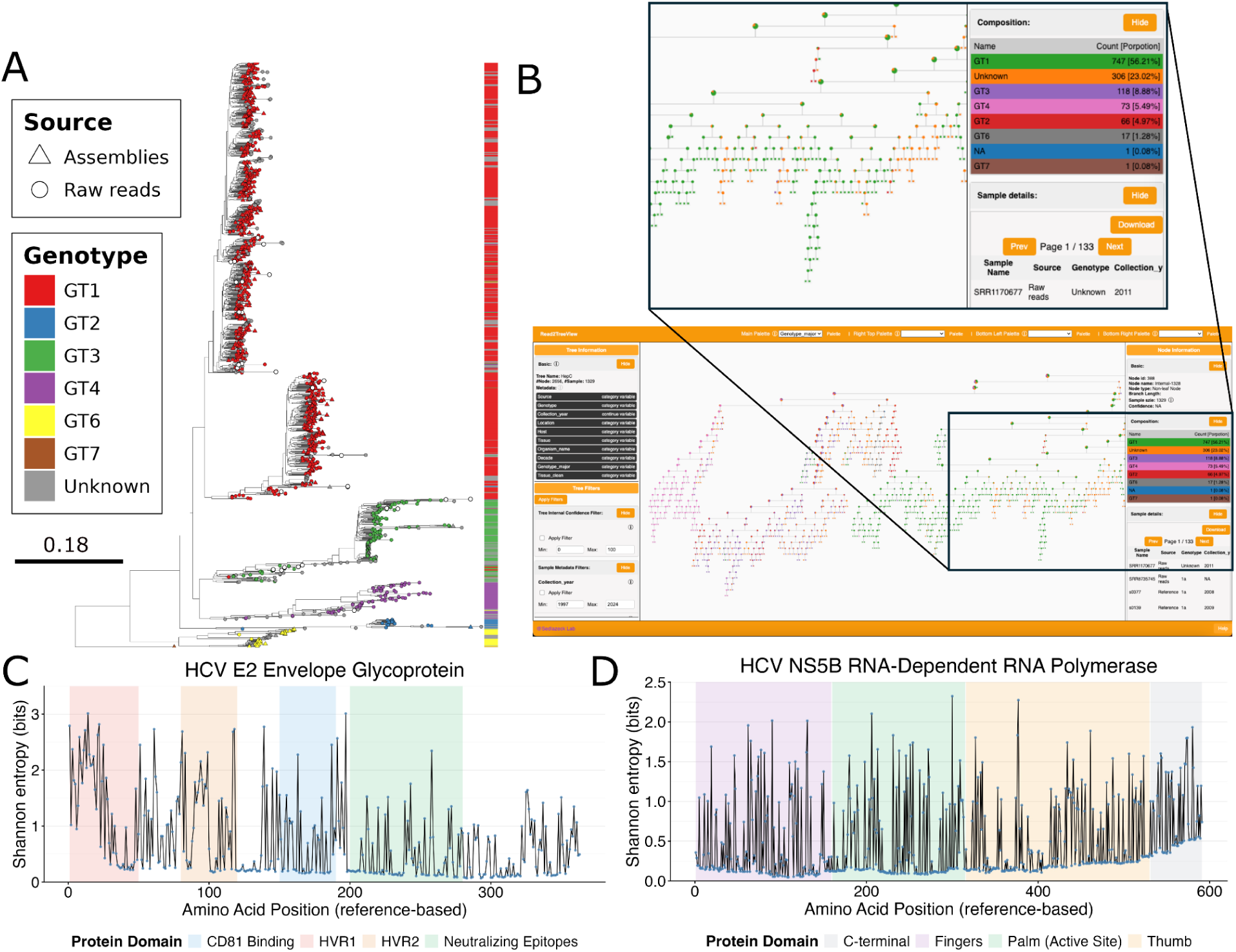
Omni2Tree phylogenomic and entropy analysis of 1,328 HCV samples. **(A)** Maximum-likelihood phylogeny inferred from 613 genome assemblies and 715 raw read datasets. Tips are colored by genotype (GT1–GT7; gray = unknown) and shaped by data source (triangles = assemblies; circles = raw reads). The color strip on the right summarizes the genotype along the tree. White tip symbols show outlier sequences. The outgroup sequence was removed for display. The scale bar represents substitutions per site. **(B)** The same tree rendered in Omni2Tree’s interactive visualization interface with genotype selected as the active metadata class, illustrating the concurrent display of multiple sample attributes that is not feasible in static tree representations (See our GitHub for the interactive HTML file https://github.com/DanielPAagustinho/omni2tree). **(C-D)** Per-position Shannon entropy across the HCV proteome for the E2 envelope glycoprotein (**C**) and the NS5B RNA-dependent RNA polymerase (**D**), calculated relative to the H77 reference strain (NC_004102) positions. Colored background regions indicate annotated functional domains. Shannon entropy is expressed in bits; a value of 0 indicates complete conservation, and the theoretical maximum for 20 amino acids is 4.32 bits.

We next examined per-position Shannon entropy across the HCV proteome using the profiles generated by Omni2Tree from all 1,328 samples. The E2 envelope glycoprotein, the principal target of neutralizing antibodies^50^, showed a sharp divide between immune-exposed and functionally constrained regions (**Figure 4C**). Positions on hypervariable regions 1 and 2 (positions 1-120) frequently exceeded 2.5 bits, with peaks at 3.0 bits, corresponding to 8-16 different amino acids at individual positions and reflecting decades of antibody-driven escape across the global HCV population^51^. The CD81 receptor-binding domain (positions 150-190) and downstream neutralizing epitope regions (positions 200-280) were far more conserved, with median entropy around 1.0 bits, indicating that receptor engagement imposes tight functional constraints even at antibody-targeted sites. This contrast helps explain why vaccines directed at the hypervariable N-terminus of E2 have failed to elicit broad protection, and points toward the more conserved receptor-binding and epitope regions as better immunogen candidates^52–54^.

The NS5B RNA-dependent RNA polymerase presented a sharply contrasting pattern (**Figure 4D**). Its palm domain active site (positions 160-315) maintained median entropy below 0.4 bits across all seven genotypes, validating it as a pan-genotypic drug target and consistent with the >95% cure rates achieved by polymerase inhibitors. The fingers domain (positions 1-159) was similarly conserved, while the thumb domain (positions 316-530) showed occasional peaks at 1.5-2.0 bits at positions that likely tolerated genotype-specific polymorphisms or resistance mutations^55^. Even the most variable NS5B positions remained below 2.5 bits, well below the E2 hypervariable peaks, confirming that polymerase function constrains evolutionary flexibility throughout the protein.

### Omni2Tree resolves hCMV phylogeny and profiles diversity across the proteome

We next applied Omni2Tree to 707 Human cytomegalovirus (hCMV) samples comprising 30 genome assemblies and 677 raw read datasets (**Supplementary tables 8** and **9**), completing the full pipeline in ∼10 days with 16 threads (4,463 CPU hours). hCMV presents a fundamentally different challenge from HCV; it is a large dsDNA herpesvirus with a ∼236 kb genome encoding over 170 open reading frames^56^. Its genome is approximately 15-fold larger than HCV’s, and its clinical dataset reflects highly heterogeneous sampling contexts spanning transplant recipients, congenitally infected neonates, and immunocompromised patients across multiple countries^57^. The resulting phylogeny (**Figure 5A**) revealed structured clustering, though interpretation is constrained by the available metadata. Unlike HCV, the hCMV dataset lacks genotype or strain annotations for the majority of samples, hCMV strains are not formally classified into discrete genotypes^58^, and most SRA accessions do not have any genotype classification. The clearest interpretable signal was the separation of laboratory-adapted and reference strains from clinical isolates, in line with convergent evolution during cell culture passage, driving recurrent mutations at loci involved in cell tropism^59^. As with HCV, assemblies and raw read datasets were distributed throughout the tree without systematic clustering by data source, confirming that the input does not introduce phylogenetic bias in Omni2Tree. The multi-metadata view generated by Omni2TreeView in **Figure 5B** illustrates why no single categorical variable organizes the hCMV tree in the way that genotype does for HCV, and its interpretive power depends on metadata quality as much as on phylogenetic resolution.

**Figure 5.**
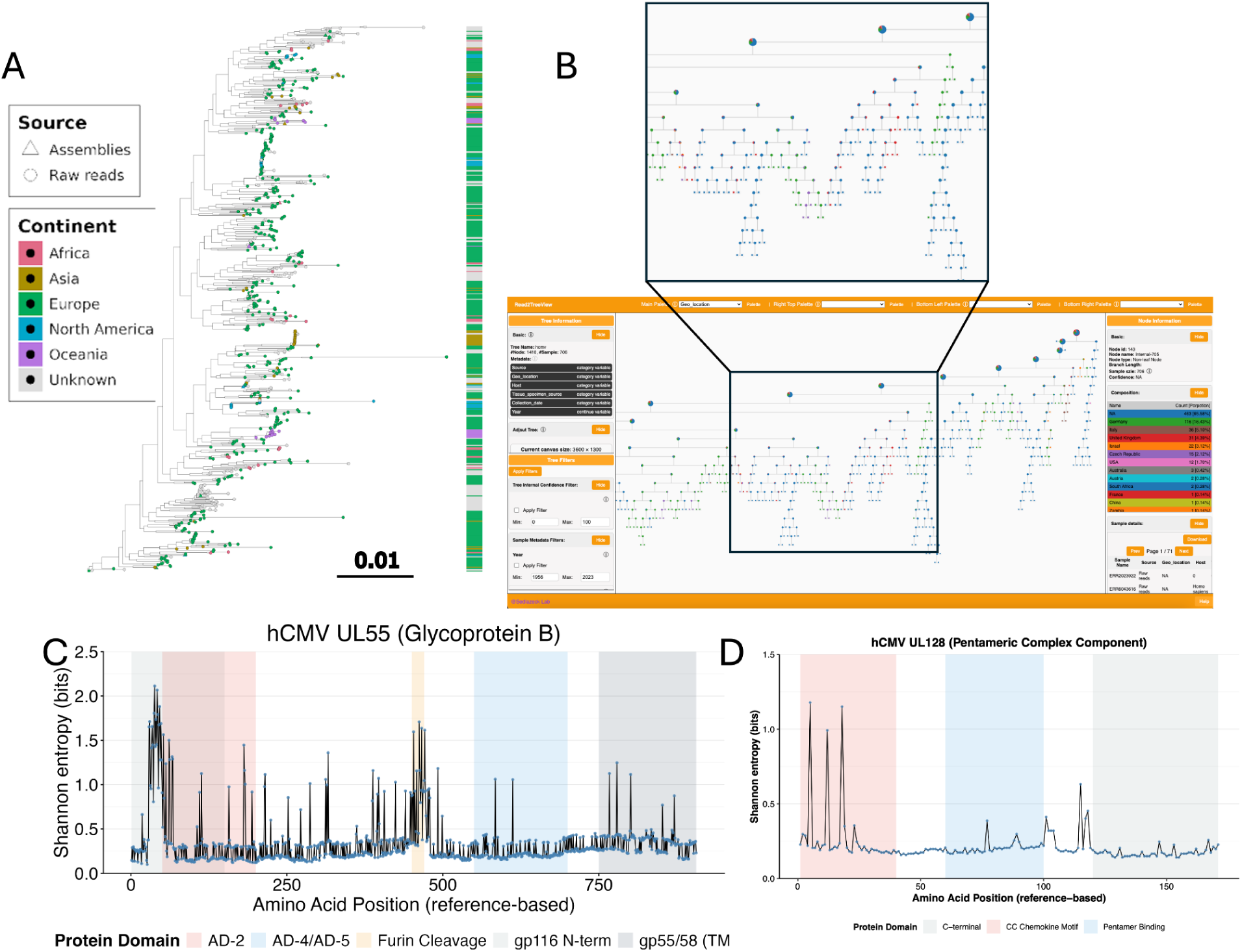
Omni2Tree phylogenomic and entropy analysis of 707 hCMV samples. (**A**) Maximum-likelihood phylogeny inferred from 30 genome assemblies and 677 raw read datasets. Tips are colored by country of origin (gray = unknown) and shaped by data source (triangles = assemblies; circles = raw reads). Murine- and macaque-derived outgroup sequences (n = 16) were removed for display. The scale bar represents substitutions per site. (**B**) The same tree shown in Omni2Tree’s interactive visualization interface, illustrating the concurrent display of multiple sample attributes. The separation of laboratory-adapted and reference strains from clinical isolates is visible without a formal genotype classifier, reflecting convergent evolution during cell culture passage. (**C–D**) Per-position Shannon entropy across the hCMV proteome for glycoprotein B (UL55; **C**) and the UL128 pentameric complex component (**D**), calculated relative to the Merlin reference strain (NC_006273) positions. Colored background regions indicate annotated functional domains. Shannon entropy is expressed in bits; a value of 0 indicates complete conservation, and the theoretical maximum for 20 amino acids is 4.32 bits.

Entropy profiles for hCMV revealed a different evolutionary regime. In glycoprotein B (UL55), an essential mediator of viral entry and cell-to-cell spread^60^, the N-terminal ectodomain (positions 1-150) carried multiple entropy peaks at 1.5-2.0 bits, consistent with immune pressure on surface-exposed residues (**Figure 5C**). The furin cleavage site (around position 460) also showed elevated entropy (∼1.7 bits), reflecting the strain-specific variation that defines gB genotypes^61^. In contrast, the AD-4/AD-5 neutralizing epitope region (positions 550-700) maintained median entropy of only 0.3-0.5 bits despite being an antibody target^62^, suggesting strong functional constraints on the fusion machinery. Current gB-based vaccine trials have shown limited efficacy^63,64^, and our entropy data suggest this may partly reflect the inclusion of the highly variable N-terminal region in immunogen design. Focusing on the conserved AD-4/AD-5 epitopes could improve cross-strain protection.

UL128, a component of the pentameric complex required for epithelial and endothelial cell entry^65^, showed a more muted entropy landscape overall (**Figure 5D**). The N-terminal CC chemokine motif (positions 1-40) was the most variable region, with several positions exceeding 1.0 bits, while the pentamer-binding region (positions 60-100) was tightly conserved (median 0.2-0.3 bits), suggesting purifying selection on the protein-protein contacts needed for pentamer assembly^66^. The lower maximum entropy in UL128 (peaks ∼1.2 bits) compared with HCV E2 (peaks >3.0 bits) likely reflects hCMV’s lower mutation rate and the different selective pressures of persistent versus acute infection.

The entropy profiles generated automatically by the Omni2Tree workflow, identify both the hypervariable regions driving immune evasion and the conserved sites that represent candidates for broadly protective vaccines and pan-genotypic antivirals.

### Omni2Tree detects co-infecting respiratory viruses in clinical metagenomic samples

Omni2Tree’s metagenomic mode extends phylogenetic reconstruction to samples containing mixed viral populations from multiple co-infecting pathogens. To validate this capability in a controlled setting, we first benchmarked the metagenomics mode against the CAMI Strain Madness challenge dataset^30^, which is a simulated complex bacterial community containing multiple co-occurring strains per species. Using a targeted 193-strain reference database, Omni2Tree recovered strain-level populations corresponding to 131 and 141 reference strains across the two CAMI datasets, with 127 strains (>90%) detected in both (**Supplementary Note**).

To assess performance on real clinical metagenomic data, we applied it to 11 metagenomic samples from two cohort studies of patients with respiratory infections during the COVID-19 pandemic (BioProjects PRJNA819439^67^ and PRJNA815970^68^; **Supplementary Table 13)**. We constructed a reference database spanning 11 respiratory viruses: SARS-CoV-2 (Wuhan-Hu-1, Delta, and Omicron variants), Influenza A, Influenza B, hMPV, HAdV-C, HAdV-F, HRV-B, HRV-C, EV-B, HCoV-NL63, and hRSV-B with three reference assemblies per virus type (**Supplementary Table 14**), and analyzed each sample without prior knowledge of which viruses were present, using Omni2Tree.

Omni2Tree partitioned reads from co-infecting viruses onto distinct phylogenetic clades within the same tree (**Figure 6A**), simultaneously placing multiple pathogens on a shared phylogeny in a single run. Note that only within-clade branch lengths are evolutionarily meaningful, while distances between clades aren’t due to sparse orthologous group overlap. For each sample, we compared our detections against the original study annotations (**Figure 6B**). In the majority of cases, Omni2Tree identified the same co-infections independently reported by the source studies, providing cross-study validation of both the detections and the original annotations (**Supplementary Table 13**). Sample 9 is among the clearest demonstrations: SARS-CoV-2 and hRSV-B reads partition cleanly onto two phylogenetically distant clades with no cross-contamination, matching the original study annotation. Sample 8 shows unambiguous FluA detection as the sole pathogen, again concordant with the source study. In samples 1 to 4, SARS-CoV-2 and HAdV-C were detected in agreement with the source annotations across all four samples. Sample 1 additionally shows concordant detection of FluB and hMPV, demonstrating that Omni2Tree correctly resolves three co-infecting viruses simultaneously from a single unassembled sample. Sample 6 is informative for sensitivity as HRV-B was present at 103,870 reads in the source study and Omni2Tree detected it correctly, placing reads within the expected rhinovirus clade. Sample 7 shows concordant detection of SARS-CoV-2 and EV-B. Samples 10 and 11 both show concordant SARS-CoV-2 detection. These concordant detections across samples from two independent cohorts sequenced with different library preparation strategies show that Omni2Tree’s marker gene approach reliably partitions reads from co-infecting viruses onto the correct phylogenetic positions without requiring prior knowledge of community composition.

**Figure 6.**
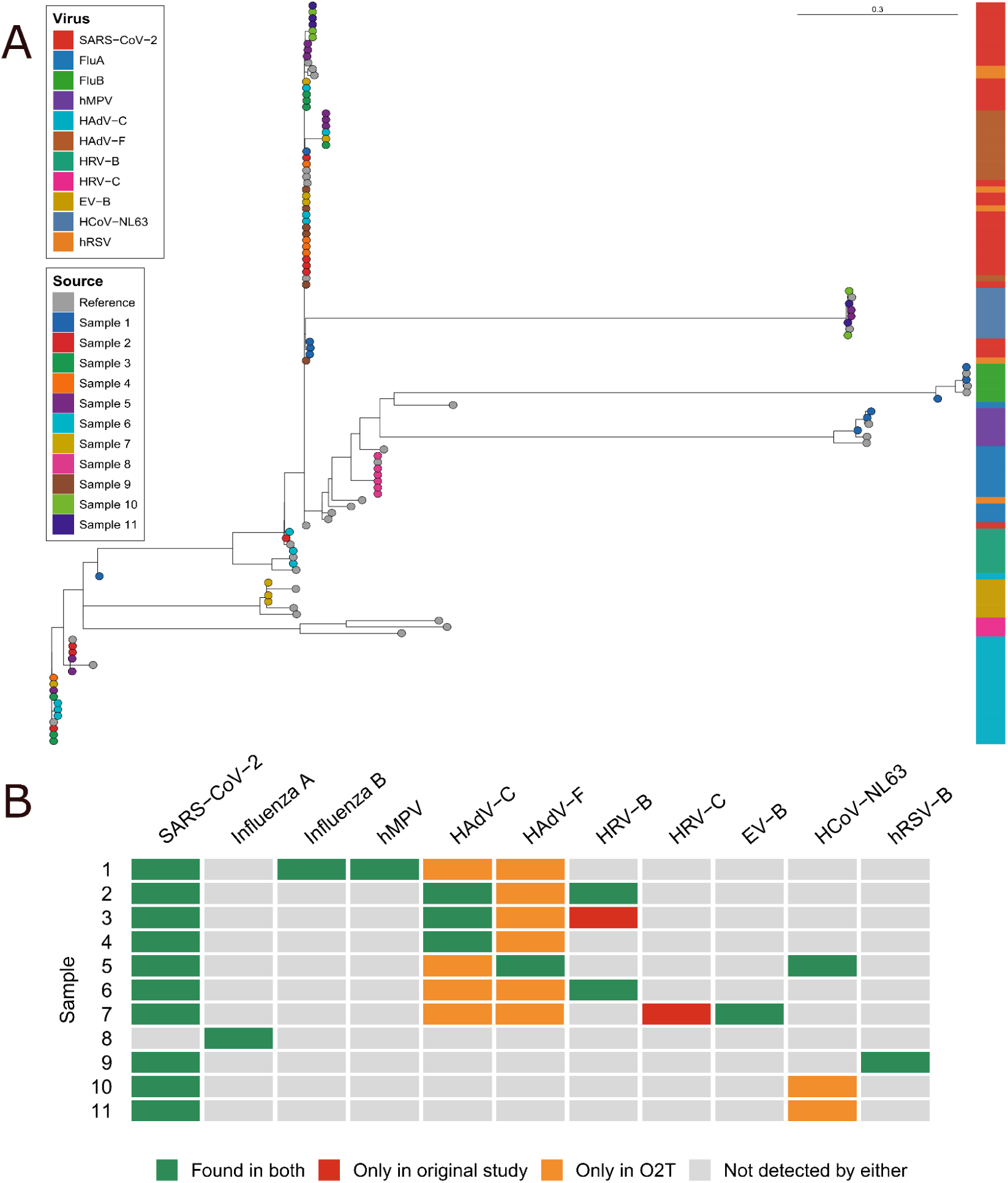
Omni2Tree resolves co-infecting respiratory viruses in clinical metagenomic samples. **(A)** Maximum-likelihood phylogeny inferred from 11 metagenomic samples spanning two cohort studies (BioProjects PRJNA819439 and PRJNA815970). Each tip represents a sequence reconstructed by Omni2Tree from reads of a single sample mapping to a given viral orthologous group; tip color indicates the sample of origin (gray = reference assemblies). The color strip on the right indicates the viral clade assignment of each tip. Reads from co-infected samples partition onto their respective viral clades without prior knowledge of community composition, demonstrating Omni2Tree’s ability to simultaneously place multiple co-infecting viruses on a shared phylogeny in a single unassembled run. Branch lengths between distinct viral clades reflect sparse inter-viral orthologous group overlap and are not interpretable as evolutionary distances; within-clade branch lengths, where shared orthologous positions exist, are phylogenetically meaningful. The scale bar represents substitutions per site. **(B)** Per-sample detection matrix across the 11 reference virus types. Green = detected and consistent with source study annotation (true positive); red = present in source study annotation but not detected by Omni2Tree; orange = detected by Omni2Tree but absent from source study annotation; gray = not expected and not detected.

Two classes of Omni2Tree-only detections were observed, both consistent with cross-alignment between closely related viruses at very low read depth rather than genuine co-infection. Across samples 1-7 (**Supplementary Table 13)**, HAdV-C and HAdV-F signals co-occurred regardless of which adenovirus type the source study reported, with signal confined to one or two marker genes with 6-30 reads in all cases except sample 5, where 617 reads (out of 16,457,244 reads) mapped to the pTP gene alone at 489x coverage but no other marker gene carried supporting signal. Given the extensive coding sequence conservation between HAdV-C and HAdV-F^69^, this pattern is in line with cross-alignment between serotypes. In samples 10 and 11, HCoV-NL63 appeared as an Omni2Tree-only detection in samples where only SARS-CoV-2 was annotated: 2 reads mapped to ORF1ab against a single NL63 reference at coverage 1 in sample 10, and 10 and 9 reads mapped against two references at coverage 5 and 4 in sample 11. Both SARS-CoV-2 and HCoV-NL63 are betacoronaviruses sharing conserved replication and structural gene sequences^70^, and the signal in both samples is confined to a single low-coverage marker gene. Omni2Tree’s metagenomic mode reports per-pathogen marker gene coverage statistics for each sample, allowing detections supported by very low read counts or confined to a single gene to be flagged as potential cross-alignment artifacts and distinguished from high-confidence detections before any clinical or epidemiological interpretation.

## Discussion

Omni2Tree removes a persistent bottleneck in viral genomics by enabling phylogenetic analysis directly from raw sequencing reads without requiring genome assembly, while simultaneously supporting mixed input types, metadata-aware visualization, and genome-wide diversity profiling. Across the datasets examined here, this design enabled rapid and consistent phylogenetic placement from short-read, long-read, and metagenomic samples, including co-infection analyses, while also generating interpretable summaries of sequence diversity through integrated entropy analysis. In doing so, Omni2Tree shifts phylogenetic analysis from a multi-step expert workflow toward a more accessible and portable framework for outbreak investigation, routine surveillance, and exploratory analysis of mixed infections from the same underlying data. This is particularly relevant when samples are heterogeneous, incomplete, or generated faster than conventional workflows can be curated.

Robustness to contamination and incomplete viral signal is not a secondary performance feature but a central requirement for practical surveillance. Viral sequencing data from clinical diagnostics, field sampling, or environmental monitoring are rarely pristine. Low viral titers, host contamination, incomplete genome recovery, and heterogeneous sequencing strategies are common features of real datasets. Across the H5N1 analyses, Omni2Tree remained robust under these conditions while distinguishing between samples that could be placed reliably and those that should be excluded because the viral signal was insufficient. This is an important property of the framework, because explicitly recognizing an insufficient signal is often more biologically and epidemiologically relevant than returning a tree position that appears confident but is only weakly supported by the data. At the same time, standard reference-guided consensus approaches introduced systematic bias from reference genome choice^18,19,71^. This bias is undetectable in routine surveillance, where only a single reference is typically selected, and it can persist even at high sequencing depth. As shown in our benchmarks, assembly-based and reference-guided mapping approaches degraded substantially under contamination and low-coverage conditions unless supplemented by additional filtering, curation, or expert intervention^14^.

An equally important aspect of Omni2Tree is that it supports a more decentralized model of viral surveillance. During recent outbreaks, centralized platforms were invaluable, but they were also necessarily selective, often prioritizing samples that met specific quality thresholds and standardized processing requirements. Omni2Tree addresses a complementary need by enabling local analysis directly from raw reads and producing a compact self-contained HTML report that can be shared with collaborators without requiring dedicated servers or specialized software. This shortens the path from sequencing to interpretation and is particularly relevant for regional laboratories, wildlife surveillance groups, and public health teams working in settings where centralized support may be limited. The same workflow also generates entropy profiles across genes and proteins, providing an immediate view of conserved and variable regions alongside the phylogeny. In the HCV and hCMV datasets, these summaries recapitulated expected contrasts between hypervariable and constrained regions, showing that Omni2Tree provides not only phylogenetic placement but also a practical first-pass view of genomic diversity relevant to immune escape, drug targeting, and vaccine design. Together, these outputs make the framework more useful for collaborative surveillance because they turn raw sequence data into interpretable results that can be shared and acted upon quickly.

The metagenomic mode extends this concept to mixed infections, where the relevant question is no longer only which virus is present, but which lineage or strain is represented and how multiple pathogens relate to one another in the same sample. In the respiratory co-infection datasets analyzed here, Omni2Tree partitioned reads from co-infecting viruses onto distinct phylogenetic clades without prior specification of sample composition, thereby adding evolutionary context to detections that would otherwise remain taxonomic labels. At the same time, our analyses also show that this mode depends strongly on the breadth and granularity of the reference database, and that weak cross-alignment between closely related viruses^69,70^ can generate low-level secondary signals. These cases emphasize that metagenomic phylogenomics should be interpreted in the context of per-gene support rather than reduced to a simple present-or-absent call.

Taken together, Omni2Tree fills a clear gap between simple classification tools and traditional phylogenomic workflows that depend on assembled genomes and centralized curation. By operating directly on raw reads, remaining robust to contamination and weak signal, and coupling phylogenetic placement with portable visualization and diversity summaries, it substantially broadens the practical scope of viral genomics. This is most relevant in exactly the settings where outbreak detection is often hardest but most important: early events, imperfect field samples, mixed infections, and regions where rapid local interpretation may be more valuable than waiting for a centralized pipeline to process only the cleanest subset of the data. In that sense, Omni2Tree does not simply speed up an existing workflow; it expands which samples, which settings, and which users can realistically participate in phylogenomic surveillance.

## Methods

### Omni2Tree workflow

Omni2Tree is organized as a modular workflow that explicitly decouples (i) the construction of a reference marker gene database from public assembly accessions, (ii) the processing and mapping of raw reads (short or long reads) against that reference database to reconstruct sequences per orthologous group, and (iii) the final combination of orthologous groups to produce a phylogeny ready for downstream analyses. In practice, this structure is implemented in three operational steps (Steps 1–3), in addition to an auxiliary script to automatically download reads from the Sequence Read Archive (SRA)^72^. Omni2Tree is available at https://github.com/DanielPAagustinho/omni2tree.

#### Step 1

A reference database is created from NCBI assemblies specified in the input CSV file. The user provides (i) an input file with taxon/strain and one or more accessions (e.g., Nucleotide accessions or assembly identifiers), and (ii) optionally an outgroups file used for phylogeny rooting and to stabilize orthology inference (avoiding generic strategies such as midpoint rooting when not appropriate). From the accessions, Step 1 retrieves the coding sequences (CDS) per taxon, prepares nucleotide and amino acid FASTAs, and performs orthologous group (OG) inference with OMA Standalone^73^. The core output of this step is (i) the marker_genes/ directory with OGs in the format required, and (ii) a dna_ref.fa file with all reference CDSs that serves as an anchor for subsequent mapping. If interrupted, the run can easily restart using the ‘--resume’ argument. In addition, auxiliary tables (e.g., mapping taxa to 5-letter codes when applicable), OMA run parameter files, and control/statistical reports on CDS and OG coverage are generated; all the generated files are organized under a working directory to enable reproducible execution and reuse of the same reference set across multiple runs.

#### Step 2

This step processes reads and maps them to the reference using minimap2^74^. It takes fastq/fastq.gz files as input and integrates them into the working directory generated in Step 1. This step supports single (default) or paired reads and allows explicit adjustment of mapping behavior (e.g., minimap2 presets suitable for ONT, PacBio or Illumina). Optionally, Step 2 can apply read deduplication (https://github.com/chanzuckerberg/czid-dedup) and downsampling^75^ controlled by target coverage and genome size. As output, Step 2 produces (i) a directory containing the consensus sequences for each sample, (ii) per-sample temporary files (original, deduplicated, and/or downsampled reads), and (iii) a tabular summary of read statistics (number of reads, average lengths, and total bases). Moreover, our design allows Step 2 to be parallelized per sample (e.g., with GNU parallel or SLURM) without rebuilding the reference database.

#### Step 3

This step first validates the metadata against both the reference taxa and the read-derived consensus sequences produced in the previous steps. Then, it combines OGs with reconstructed sequences and runs IQ-TREE3^76^ to infer a maximum-likelihood phylogenetic tree. After that, Step 3 performs a Shannon entropy analysis on the merged alignments by converting orthogroup MSAs into position-level tables, calculating entropy across all genes or protein sequences, and producing genome-wide and per-gene plots, with optional annotation of known protein domains. Finally, metadata and tree labels are integrated to generate an interactive HTML visualization with Omni2TreeView. This phase consolidates the multi-gene evidence and produces a final tree that can be used directly for diversity analysis, surveillance, and/or comparison between samples derived from raw reads and assembled genomes within the same framework.

#### Omni2TreeSRA

The Omni2TreeSRA script supports batch download from SRA using the SRA Toolkit: from a file with SRA identifiers, it queries the metadata with *esearch/efetch*, downloads each RUN/EXPERIMENT, and converts it to FASTQ files. The output files are renamed with the taxon/strain identifier, and a report summarizing the downloads is generated.

With these three steps and the download script, Omni2Tree integrates reference construction, read processing, and tree inference into a reproducible workflow.

### Omni2Tree for Metagenomics

The input to Omni2Tree-metagenomics is the same as Omni2Tree for viruses, including reference samples and the sequencing reads. The difference is that the sequencing reads come from environmental samples in FASTQ format. Omni2Tree follows the steps of Read2Tree with some modifications, since Read2Tree was designed to infer phylogeny when all reads belong to one species in a sample in FASTQ format. However, in a metagenomic sample, there are reads from different species in a FASTQ file.

The first step in Omni2Tree-metagenomics is that the OGs are aligned using the MAFFT multiple sequence aligner^77^. Then, the sequencing reads are aligned to reference gene markers (in nucleotides) using the minimap2^74^ aligner (for either short or long reads). Then, reads that are mapped to each gene from the same species are collected. For those input strains that have enough reads mapped (default 2 reads per gene), a consensus sequence is generated. This step can be done in parallel for all input read samples. Next, each consensus sequence that is generated for each of the reference strains (if any) is added to the multiple sequence alignment (MSA). Finally, the MSA is used with IQtree to infer the phylogenetic tree.

### Benchmarking Omni2Tree accuracy and performance

#### Short-read benchmark (H5N1)

Raw paired-end Illumina reads for 20 H5N1 samples from the 2024 US dairy cattle outbreak (clade 2.3.4.4b; BioProject PRJNA1102327) were downloaded using fasterq-dump (SRA Toolkit v3.0.10) and compressed with gzip. Median sequencing depth across samples was 3,155x. An additional sample from clade 2.3.2.1c (SRR11611114) was included to serve as a cross-clade accuracy control. This sample was processed identically to the outbreak samples and was also included as a GenBank assembly in the reference panel (see below). Read depth and taxonomic composition for all 21 samples were assessed with Kraken2 against a custom database, and the results are reported in **Supplementary Table 1**.

Eleven H5N1 reference assemblies spanning 1996–2024 across multiple hosts and geographic origins were included as phylogenetic context in all approaches (**Supplementary Table 2**). Reads were downsampled to 300x, 100x, 30x, 10x, 5x, 2x, 1x, and 0.5x nominal coverage using Rasusa v0.7.0 with a target genome size of 13.6 kb. For the contamination experiment, bovine genomic reads (SRR5753525; Illumina NextSeq 500, 150 bp paired-end) were spiked into three samples with the highest viral read fractions (SRR28912806, SRR28912809, SRR28912812; 91–93% Influenza A) at contamination levels of 25%, 50%, 75%, and 90% of total reads. Each contamination level was built cumulatively; the 50% condition was produced by adding reads to the 25% sample, and so on, to ensure consistency of the contaminating read set across levels.

Four phylogenetic approaches were compared. For Approach 1, reads were assembled with MegaHit v1.2.9 (default parameters, minimum contig length 200 bp). Contigs ≥500 bp were retained and concatenated with 50-N spacers to produce a single sequence per sample, which was necessary to avoid multiple tips per sample in the final tree. Concatenated assemblies and the 11 reference sequences were aligned with MAFFT v7 (--auto), and trees were inferred with IQ-TREE v2.2 (GTR+G, 1,000 ultrafast bootstrap replicates). For Approach 2, reads were mapped with BWA-MEM to two clade 2.3.4.4b reference genomes: a closely related cattle isolate (A/cattle/Texas/56283/2024) and a divergent avian isolate (A/bald eagle/Florida/W22-134-OP/2022). Consensus sequences were generated with bcftools v1.9 mpileup and call, aligned with MAFFT, and trees were inferred with IQ-TREE as above. Both reference conditions were run to expose reference-induced placement bias, which is undetectable when only a single reference is used. For Approach 3, trees were constructed directly from reads using MashTree (k-mer size 21, sketch size 10,000, mindepth 0), with both reads and reference FASTAs provided as input. For Approach 4, raw reads were processed with Omni2Tree using default parameters and a reference database constructed from the 11 reference strains. IQ-TREE v2.2 was run on the concatenated amino acid alignment output by Omni2Tree (LG+G, 1,000 ultrafast bootstrap replicates); LG+G rather than GTR+G was used because Omni2Tree marker gene alignments are amino acid sequences.

Phylogenetic placement accuracy for the cross-clade control sample (SRR11611114) was quantified as the patristic distance between the raw-read tip and the cognate GenBank assembly tip within the same tree, computed from the cophenetic distance matrix of the inferred phylogeny. A distance of zero indicates perfect co-placement with the reference assembly. For the contamination experiment, patristic distance was computed between each contaminated version of a sample and its uncontaminated baseline tip within a single combined tree inferred from all contamination levels simultaneously, with tip labels encoding both sample identity and contamination level. All patristic distances were computed using the ape package v5.x in R. Topological stability was assessed by computing Robinson-Foulds distances between each subsampled tree and the corresponding full-coverage tree for the same approach, restricted to outbreak sample tips. Robinson-Foulds distances were normalized by the maximum possible RF distance for the given number of tips (2n−6 for unrooted trees with n tips) and converted to topological accuracy as 1 − RF_normalized. Bootstrap support values ≤70 were collapsed to polytomies prior to all topological comparisons. All post-analysis visualization was performed in R using ggplot2. Wall-clock time was recorded from step-level bash timestamps within the pipeline scripts; CPU time was captured via SLURM sacct job accounting. All jobs ran on a cluster with 16 CPUs and 32 GB RAM per job.

### Long-read benchmark (RSV)

Twenty-two RSV/A samples from BioProject PRJNA980575 sequenced with ONT were retrieved using fasterq-dump. Read quality was assessed with NanoPlot v1.40.0; mean read lengths ranged from 300–700 bp, as expected from amplicon-based sequencing protocols common in respiratory virus surveillance. Per-sample coverage was variable and generally low.

Two phylogenetic approaches were compared. For reference-guided consensus, reads were mapped with minimap2 v2.24 (--ax map-ont) to an RSV/A consensus sequence from 53 2014 RSV/A assemblies^78^ as primary reference and to HRSV/A/England/397/2017 (NCBI accession PP109421.1; GISAID EPI_ISL_412866) as a secondary reference for samples failing initial mapping. Consensus sequences were generated using bcftools v1.15 mpileup and call (--ploidy 1, -d 10000), aligned with MAFFT v7, and trees were inferred with IQ-TREE v2.2 (GTR+G, 1,000 ultrafast bootstrap replicates). Standard long-read assembly tools (Flye v2.9.1, Canu v2.2) were used but were not viable for this data due to insufficient read length and coverage. For Omni2Tree, the same raw reads were processed directly without any assembly or reference-guided steps, using a database constructed from a panel of RSV/A and RSV/B reference strains (**Supplementary Table 3**).

### Phylogenomic analysis of HCV and hCMV samples using Omni2Tree

We retrieved all available hepatitis C virus (HCV) and human cytomegalovirus (hCMV) sequencing data from public repositories through March 2025. For HCV, we obtained 1,328 samples comprising 613 assemblies from GenBank and 715 raw read datasets from the NCBI SRA. For hCMV, we collected 707 samples, including 30 assemblies and 677 raw read datasets. Genotype information for HCV samples was extracted directly from existing SRA and GenBank metadata annotations, which indicated representation across genotypes 1-7. For hCMV, metadata were notably sparse, with limited strain or lineage information available for most samples. We then executed the 3 modular steps of Omni2Tree to generate a high-quality phylogenomic analysis. This approach enabled us to process both genome assemblies and raw sequencing reads in a unified workflow. Phylogenetic trees were constructed using IQ-TREE v2.2 with automatic model selection and 1,000 ultrafast bootstrap replicates.

Association between sample metadata and phylogenetic clustering was quantified using Cramér’s V^79^, a chi-squared-based statistic normalized to the range [0, 1], where 0 indicates no association, and 1 indicates perfect association between two categorical variables.

### Reference-based Shannon entropy calculation

To ensure biologically meaningful position numbering, we implemented reference-based coordinate mapping for entropy calculations. All alignment positions were mapped to well-characterized reference strains: H77 (NC_004102) for HCV and Merlin (NC_006273) for hCMV. We analyzed therapeutically relevant genes: for HCV, Core, E1, E2, NS3, NS4A, NS4B, NS5A, and NS5B; for hCMV, UL55, UL75, UL115, UL128, UL130, UL83, and UL122/123. For each gene alignment, we identified the reference sequence and created a mapping from each alignment column to the corresponding ungapped position in the reference. Alignment columns where the reference contained gaps were excluded from analysis. This approach ensures that reported positions correspond to actual amino acid positions in the reference protein, enabling direct application of structural and functional annotations from literature.

Shannon entropy was calculated per reference amino acid position as described previously^78^, with values ranging from 0 bits (complete conservation) to 4.32 bits (maximum observed diversity). Positions were filtered to include only those with at least 5 sequences. All analyses were performed using custom Python and R scripts available at the Omni2Tree GitHub repository.

### Metagenomics Benchmarking

We used the CAMI^30^ StrainMadness benchmark dataset to evaluate the performance of Omni2Tree in metagenomics mode. We use the output tree from Omni2Tree and report the content of the metagenomics sample at strain and species levels. We also use the read mapping (intermediate output of Omni2Tree) to classify each read. Even though Omni2Tree is not designed for classification task, we compared it with Kraken^28^ version 2.1.6 using the Standard-16 index k2_standard_16_GB_20250714.tar.gz from https://benlangmead.github.io/aws-indexes. We calculated the classification accuracy at the lowest taxonomy level of the CAMI true set (species) and followed the definition of false positive (FP, read classified to an incorrect taxon id) and true positive (TP, read classified to the correct taxon id) as literature^28^. We define precision as TP/(TP + FP) and recall as TP/P. F1 is defined as 2*precision*recall/(precision+recall) (**Supplementary Note 2**).

The co-infection analysis was done using Omni2Tree in metagenomics mode. We downloaded the 11 sequencing read datasets automatically as part of Omni2Tree (**Supplementary Table 13**) and generated the reference gene markers using OMA standalone^73^ (**Supplementary Table 14**). The phylogenetic tree is inferred using IQ-TREE2^80^. We used the co-infection reported in each study visualized in **Figure 6A-B**.

### Omni2TreeView

We developed an interactive visualization framework, Omni2TreeView, to explore the tree structures generated by the Omni2Tree main program. Omni2Treeview leverages D3.js (https://github.com/d3/d3) to render tree structures and associated metadata, with metadata explicitly displayed for all leaf nodes. To support the integration of multiple metadata categories and enable tree rendering conditioned on these annotations, Omni2TreeView incorporates Vue.js (https://vuejs.org/) to construct a user interface layered on top of the rendered tree.

The tree view and the user interface are tightly coupled and fully interactive, allowing users to dynamically filter the tree based on any metadata category. In addition, Omni2TreeView enables inspection of node-level details through direct interaction with individual tree nodes. While metadata are displayed directly for leaf nodes, values associated with internal nodes are computed through the aggregation of their descendant leaves.

Omni2Tree is implemented as a lightweight Python script together with an HTML template, with minimal external dependencies; Biopython is the only required third-party library. This design facilitates straightforward customization of metadata and enables users to easily regenerate the interactive tree visualization. Omni2TreeView is available at the ‘view’ subfolder in the Omni2Tree repository https://github.com/DanielPAagustinho/omni2tree.

## Data Availability

All raw sequencing data used in this study are publicly available. H5N1 benchmark samples are deposited under BioProject PRJNA1102327. RSV/A long-read samples are available under BioProject PRJNA980575. HCV and hCMV raw reads and assemblies were retrieved from the NCBI Sequence Read Archive and GenBank through March 2025; individual accession numbers are listed in Supplementary Tables 1-5. Metagenomic co-infection samples are available under BioProjects PRJNA819439 and PRJNA815970. CAMI Strain Madness benchmark datasets are available at https://cami-challenge.org/datasets/. H5N1 outbreak samples from the Webby group are being deposited on NCBI, which will be reported here during revision. Reference assemblies used for all analyses are listed in Supplementary Tables 2, 4, 5, 7, and 13.

## Code Availability

Omni2Tree is freely available at https://github.com/DanielPAagustinho/omni2tree. Omni2TreeView, as well as all analysis scripts used for benchmarking, visualization, and entropy calculation are available at the same repository.

## Authors contributions

S.M. contributed to analysis, tool development, and writing. A.C. contributed to analysis, tool development, and writing. X.Z. contributed to tool development and writing. R.J.W. contributed data. A.S.B. and R.L.P. contributed samples. N.M.N. contributed to sample collection and metadata. F.J.S. supervised the project and contributed to planning and writing. D.P.A. conceived the study and contributed to planning, analysis, tool development, and writing.

## Reference

1. Günthard, H. F., Kusejko, K. & Kouyos, R. D. Phylogenetics and molecular evolution to understand and curb the HIV pandemic. Nat. Rev. Microbiol. 24, 76–92 (2026).

2. Li, T. et al. Phylogenetic supertree reveals detailed evolution of SARS-CoV-2. Sci. Rep. 10, 22366 (2020).

3. Attwood, S. W., Hill, S. C., Aanensen, D. M., Connor, T. R. & Pybus, O. G. Phylogenetic and phylodynamic approaches to understanding and combating the early SARS-CoV-2 pandemic. Nat. Rev. Genet. 23, 547–562 (2022).

4. Nguyen, T.-Q., et al. Emergence and interstate spread of highly pathogenic avian influenza A(H5N1) in dairy cattle. bioRxiv (2024) doi:10.1101/2024.05.01.591751.

5. Damodaran, L., Jaeger, A. S. & Moncla, L. H. Ecology and spread of the North American H5N1 epizootic. Nature 649, 432–441 (2026).

6. Havens, J. L. et al. Dynamics of natural selection preceding human viral epidemics and pandemics. Cell (2026) doi:10.1016/j.cell.2026.02.006.

7. Dylus, D., Altenhoff, A., Majidian, S., Sedlazeck, F. J. & Dessimoz, C. Inference of phylogenetic trees directly from raw sequencing reads using Read2Tree. Nat. Biotechnol. 42, 139–147 (2024).

8. Inferring phylogenies from pandemic-scale genome datasets. Nat. Genet. 55, 734–735 (2023).

9. De Maio, N. et al. Maximum likelihood pandemic-scale phylogenetics. Nat. Genet. 55, 746–752 (2023).

10. Hadfield, J. et al. Nextstrain: real-time tracking of pathogen evolution. Bioinformatics 34, 4121–4123 (2018).

11. Raghavan, V., Kraft, L., Mesny, F. & Rigerte, L. A simple guide to de novo transcriptome assembly and annotation. Brief. Bioinform. 23, (2022).

12. Atxaerandio-Landa, A. et al. A practical bioinformatics workflow for routine analysis of bacterial WGS data. Microorganisms 10, 2364 (2022).

13. Nguinkal, J. A. et al. Assessment of the pathogen genomics landscape highlights disparities and challenges for effective AMR Surveillance and outbreak response in the East African community. BMC Public Health 24, 1500 (2024).

14. Jansz, N. & Faulkner, G. J. Viral genome sequencing methods: benefits and pitfalls of current approaches. Biochem. Soc. Trans. 52, 1431–1447 (2024).

15. Castro, C. J., Marine, R. L., Ramos, E. & Ng, T. F. F. The effect of variant interference on de novo assembly for viral deep sequencing. BMC Genomics 21, 421 (2020).

16. Han, S. M., Kubo, Y., Robert, A., Baguelin, M. & Ariyoshi, K. Impact of viral co-detection on the within-host viral diversity of influenza patients. Viruses 17, 152 (2025).

17. Wagner, D. D. et al. VPipe: An automated bioinformatics platform for assembly and management of viral next-generation sequencing data. Microbiol. Spectr. 10, e0256421 (2022).

18. Norri, T. & Mäkinen, V. Tackling reference bias in genotyping by using founder sequences with PanVC 3. Bioinform. Adv. 4, vbae027 (2024).

19. Maurstad, M. F. et al. Reference genome bias in light of species-specific chromosomal reorganization and translocations. Genome Biol. 26, 355 (2025).

20. Ondov, B. D. et al. Mash: fast genome and metagenome distance estimation using MinHash. Genome Biol. 17, 132 (2016).

21. Katz, L. S. et al. Mashtree: a rapid comparison of whole genome sequence files. J. Open Source Softw. 4, 1762 (2019).

22. Ondov, B. D. et al. Mash Screen: high-throughput sequence containment estimation for genome discovery. Genome Biol. 20, 232 (2019).

23. Georgakopoulou, V. E. Insights from respiratory virus co-infections. World J. Virol. 13, 98600 (2024).

24. Nickbakhsh, S. et al. Virus-virus interactions impact the population dynamics of influenza and the common cold. Proc. Natl. Acad. Sci. U. S. A. 116, 27142–27150 (2019).

25. Agustinho, D. P. et al. Unveiling microbial diversity: harnessing long-read sequencing technology. Nat. Methods 21, 954–966 (2024).

26. Nurk, S., Meleshko, D., Korobeynikov, A. & Pevzner, P. A. metaSPAdes: a new versatile metagenomic assembler. Genome Res. 27, 824–834 (2017).

27. Kang, D. D. et al. MetaBAT 2: an adaptive binning algorithm for robust and efficient genome reconstruction from metagenome assemblies. PeerJ 7, e7359 (2019).

28. Wood, D. E., Lu, J. & Langmead, B. Improved metagenomic analysis with Kraken 2. Genome Biol. 20, 257 (2019).

29. Nguyen, M. & Schatz, M. Perseus: Lineage-aware refinement of Kraken2 taxonomic classification for long read metagenomes. bioRxiv (2026) doi:10.64898/2026.03.06.710148.

30. Meyer, F. et al. Critical Assessment of Metagenome Interpretation: the second round of challenges. Nat. Methods 19, 429–440 (2022).

31. Shen, C., Wedell, E., Pop, M. & Warnow, T. TIPP3 and TIPP3-fast: Improved abundance profiling in metagenomics. PLOS Computational Biology 21, e1012593 (2025).

32. Liu, R. et al. Constructing phylogenetic trees for microbiome data analysis: A mini-review. Comput. Struct. Biotechnol. J. 23, 3859–3868 (2024).

33. Bouckaert, R. et al. BEAST 2.5: An advanced software platform for Bayesian evolutionary analysis. PLoS Comput. Biol. 15, e1006650 (2019).

34. Gao, Y. et al. Benchmarking short-read metagenomics tools for removing host contamination. Gigascience 14, (2025).

35. Li, H. Aligning sequence reads, clone sequences and assembly contigs with BWA-MEM. arXiv [q-bio.GN] (2013).

36. Danecek, P. et al. Twelve years of SAMtools and BCFtools. Gigascience 10, (2021).

37. Gómez-Del Rosario, A., et al. A bench-to-data analysis workflow for respiratory syncytial virus whole-genome sequencing with short and long-read approaches. Genome Med. 18, 9 (2026).

38. Kolmogorov, M., Yuan, J., Lin, Y. & Pevzner, P. A. Assembly of long, error-prone reads using repeat graphs. Nat. Biotechnol. 37, 540–546 (2019).

39. Koren, S. et al. Canu: scalable and accurate long-read assembly via adaptive*k*-mer weighting and repeat separation. Genome Res. 27, 722–736 (2017).

40. Min, J.-Y. et al. Mammalian adaptation in the PB2 gene of avian H5N1 influenza virus. J. Virol. 87, 10884–10888 (2013).

41. Zhang, L. et al. Emergence of mammalian-adaptive PB2 mutations enhances polymerase activity and pathogenicity of cattle-derived H5N1 influenza A virus. Nat. Commun. 17, 1011 (2025).

42. Echeverría, N., Moratorio, G., Cristina, J. & Moreno, P. Hepatitis C virus genetic variability and evolution. World J. Hepatol. 7, 831–845 (2015).

43. De Luca, A. et al. Two distinct hepatitis C virus genotype 1a clades have different geographical distribution and association with natural resistance to NS3 protease inhibitors. Open Forum Infect. Dis. 2, ofv043 (2015).

44. Pérez, A. B. et al. Increasing importance of European lineages in seeding the hepatitis C virus subtype 1a epidemic in Spain. Euro Surveill. 24, (2019).

45. Newman, R. M. et al. Whole genome pyrosequencing of rare hepatitis C virus genotypes enhances subtype classification and identification of naturally occurring drug resistance variants. J. Infect. Dis. 208, 17–31 (2013).

46. Bonsall, D. et al. ve-SEQ: Robust, unbiased enrichment for streamlined detection and whole-genome sequencing of HCV and other highly diverse pathogens. F1000Res. 4, 1062 (2015).

47. Perpiñán, E. et al. Hepatitis C virus early kinetics and resistance-associated substitution dynamics during antiviral therapy with direct-acting antivirals. J. Viral Hepat. 25, 1515–1525 (2018).

48. Singer, J. B. et al. Interpreting viral deep sequencing data with GLUE. Viruses 11, 323 (2019).

49. Depledge, D. P. et al. Specific capture and whole-genome sequencing of viruses from clinical samples. PLoS One 6, e27805 (2011).

50. Tzarum, N., Wilson, I. A. & Law, M. The neutralizing face of hepatitis C virus E2 envelope glycoprotein. Front. Immunol. 9, 1315 (2018).

51. Sevvana, M., Keck, Z., Foung, S. K. & Kuhn, R. J. Structural perspectives on HCV humoral immune evasion mechanisms. Curr. Opin. Virol. 49, 92–101 (2021).

52. Guest, J. D. & Pierce, B. G. Structure-based and rational design of a hepatitis C virus vaccine. Viruses 13, 837 (2021).

53. Velázquez-Moctezuma, R. et al. Mechanisms of hepatitis C virus escape from vaccine-relevant neutralizing antibodies. Vaccines (Basel) 9, 291 (2021).

54. Skinner, N. E. Broadly neutralizing antibody characteristics in hepatitis C virus infection and implications for vaccine design. Vaccines (Basel) 13, 612 (2025).

55. Boyce, S. E. et al. Structural and regulatory elements of HCV NS5B polymerase--β-loop and C-terminal tail--are required for activity of allosteric thumb site II inhibitors. PLoS One 9, e84808 (2014).

56. Grgic, I. & Gorenec, L. Human Cytomegalovirus (HCMV) genetic diversity, drug resistance testing and prevalence of the resistance mutations: A literature review. Trop. Med. Infect. Dis. 9, 49 (2024).

57. Charles, O. J. et al. Genomic and geographical structure of human cytomegalovirus. Proc. Natl. Acad. Sci. U. S. A. 120, e2221797120 (2023).

58. Peterson, C. S., Bailey, I. T., Lanchy, J.-M., Gallego, I. & Ryckman, B. J. Interstrain recombinants of human cytomegalovirus uncouple glycoprotein display, virion infectivity, and spread characteristics. J. Virol. 100, e0159225 (2026).

59. Dargan, D. J. et al. Sequential mutations associated with adaptation of human cytomegalovirus to growth in cell culture. J. Gen. Virol. 91, 1535–1546 (2010).

60. Tang, J., Frascaroli, G., Lebbink, R. J., Ostermann, E. & Brune, W. Human cytomegalovirus glycoprotein B variants affect viral entry, cell fusion, and genome stability. Proc. Natl. Acad. Sci. U. S. A. 116, 18021–18030 (2019).

61. Stangherlin, L. M. et al. Positively selected sites at HCMV gB furin processing region and their effects in cleavage efficiency. Front. Microbiol. 8, 934 (2017).

62. Wiegers, A.-K., Sticht, H., Winkler, T. H., Britt, W. J. & Mach, M. Identification of a neutralizing epitope within antigenic domain 5 of glycoprotein B of human cytomegalovirus. J. Virol. 89, 361–372 (2015).

63. Nelson, C. S. et al. HCMV glycoprotein B subunit vaccine efficacy mediated by nonneutralizing antibody effector functions. Proc. Natl. Acad. Sci. U. S. A. 115, 6267–6272 (2018).

64. Gomes, A. C., Griffiths, P. D. & Reeves, M. B. The humoral immune response against the gB vaccine: Lessons learnt from protection in solid organ transplantation. Vaccines (Basel) 7, 67 (2019).

65. Straschewski, S. et al. Protein pUL128 of human cytomegalovirus is necessary for monocyte infection and blocking of migration. J. Virol. 85, 5150–5158 (2011).

66. Kschonsak, M. et al. Structural basis for HCMV Pentamer receptor recognition and antibody neutralization. Sci. Adv. 8, eabm2536 (2022).

67. Iša, P. et al. Metagenomic analysis reveals differences in the co-occurrence and abundance of viral species in SARS-CoV-2 patients with different severity of disease. BMC Infect. Dis. 22, 792 (2022).

68. Ramos, N. et al. A multiplex-NGS approach to identifying respiratory RNA viruses during the COVID-19 pandemic. Arch. Virol. 168, 87 (2023).

69. Wang, Y. et al. Immunological study of reconstructed common ancestral sequence of Adenovirus hexon protein. Front. Microbiol. 12, 717047 (2021).

70. Castillo, G. et al. Molecular mechanisms of human coronavirus NL63 infection and replication. Virus Res. 327, 199078 (2023).

71. Valiente-Mullor, C. et al. One is not enough: On the effects of reference genome for the mapping and subsequent analyses of short-reads. PLoS Comput. Biol. 17, e1008678 (2021).

72. Katz, K. et al. The Sequence Read Archive: a decade more of explosive growth. Nucleic Acids Res. 50, D387–D390 (2022).

73. Altenhoff, A. M. et al. OMA standalone: orthology inference among public and custom genomes and transcriptomes. Genome Res. 29, 1152–1163 (2019).

74. Li, H. Minimap2: pairwise alignment for nucleotide sequences. Bioinformatics 34, 3094–3100 (2018).

75. Hall, M. Rasusa: Randomly subsample sequencing reads to a specified coverage. J. Open Source Softw. 7, 3941 (2022).

76. Wong, T. et al. IQ-TREE 3: Phylogenomic inference software using complex evolutionary models. EcoEvoRxiv (2025) doi:10.32942/x2p62n.

77. Katoh, K. & Standley, D. M. MAFFT multiple sequence alignment software version 7: improvements in performance and usability. Mol. Biol. Evol. 30, 772–780 (2013).

78. Avadhanula, V. et al. Evolutionary dynamics of Respiratory Syncytial Virus in pre-pandemic, pandemic, and post-pandemic periods in Houston, Texas, USA. bioRxiv (2025) doi:10.1101/2025.08.06.668939.

79. Cramér, H. Mathematical Methods of Statistics. in The foundational source (Princeton University Press, 1946).

80. Minh, B. Q. et al. IQ-TREE 2: New models and efficient methods for phylogenetic inference in the genomic era. Mol. Biol. Evol. 37, 1530–1534 (2020).

