## Supplementary Figures for "Rapid phylogenomic analysis for viral surveillance and metagenomic profiling with Omni2Tree"

#### Supplementary Materials for Omni2Tree study

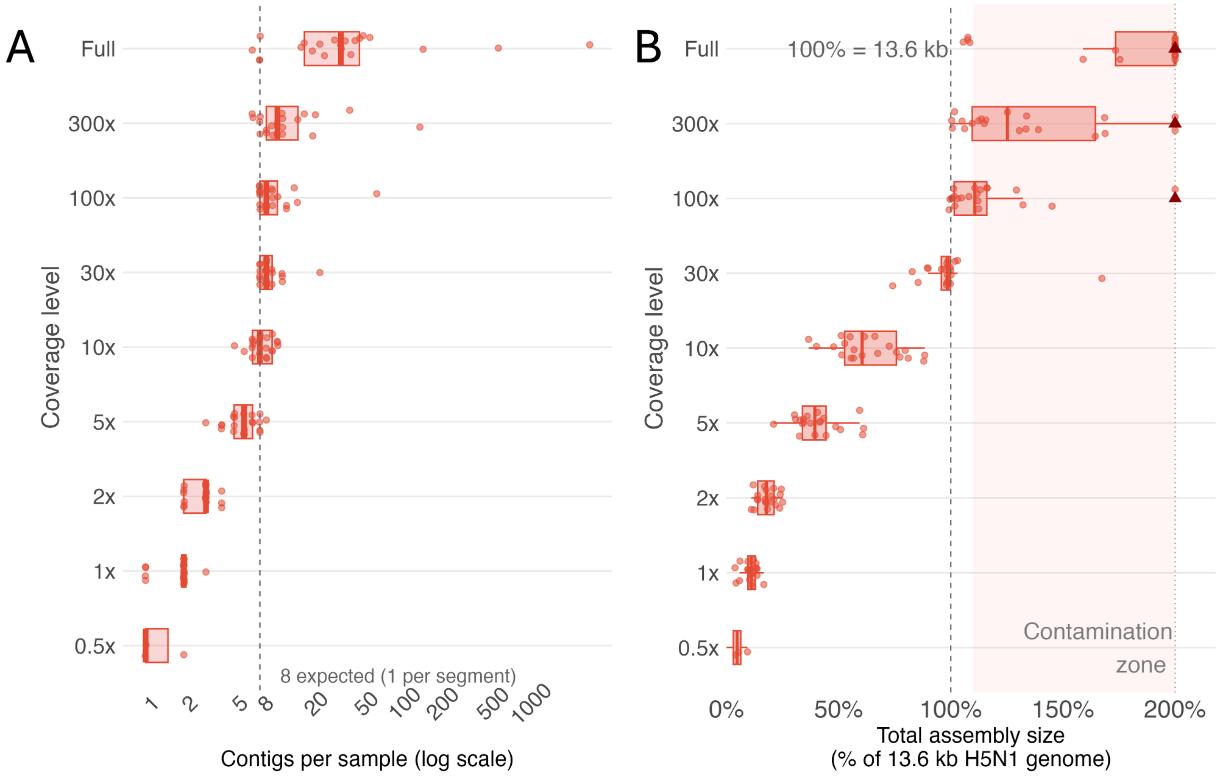


**Supplementary Figure 1. (A) Contig count per sample on a log scale**, after filtering contigs shorter than 500 bp. The dashed line indicates the expected 8 contigs (one per H5N1 genome segment). Failed assemblies excluded. **(B) Total assembly size per sample as a percentage of the expected 13.6 kb H5N1 reference genome.** Values exceeding 100% likely indicate host DNA contamination in the assembly. Axis capped at 200%; outliers shown as labeled triangles with true values.

**Supplementary Figure 2-6. Phylogenetic trees for all approaches across coverage levels.** Trees were inferred from 21 H5N1 short-read samples and 11 reference assemblies at six coverage levels (full, 300x, 100x, 30x, 10x, 5x, 2x, 1x, 0.5x). Each panel shows the tree produced by the indicated approach at the indicated coverage level, rooted on A/goose/Guangdong/1/1996 as the ancestral outgroup. Tip colors indicate sample category: gray circles = cattle outbreak samples (n=20, clade 2.3.4.4b); red circles = SRR11611114 (clade 2.3.2.1c raw reads); green circles = A/flamingo/Kazakhstan/6570/2015 (cognate assembly for SRR11611114); purple circles = A/bald eagle/Florida/W22-134-OP/2022; blue circles = A/cattle/Texas/56283/2024; light gray circles = remaining reference assemblies. Only the flamingo, bald eagle, and cattle Texas assembly tips are labeled to highlight reference positions relevant to approach-specific biases. Bootstrap support ≤70 was collapsed prior to visualization for IQ-TREE-based approaches (MegaHit + IQ-TREE, Alignment + Consensus, Omni2Tree + IQ-TREE); MashTree produces unrooted neighbor-joining trees without bootstrap support. "Tree generation failed" indicates that insufficient reads were available to complete the assembly or alignment step at that coverage level. "SRR11611114 absent" indicates that the sample was excluded from the Omni2Tree tree due to insufficient H5N1 read depth after downsampling, reflecting the method's transparent quality threshold rather than a pipeline failure.

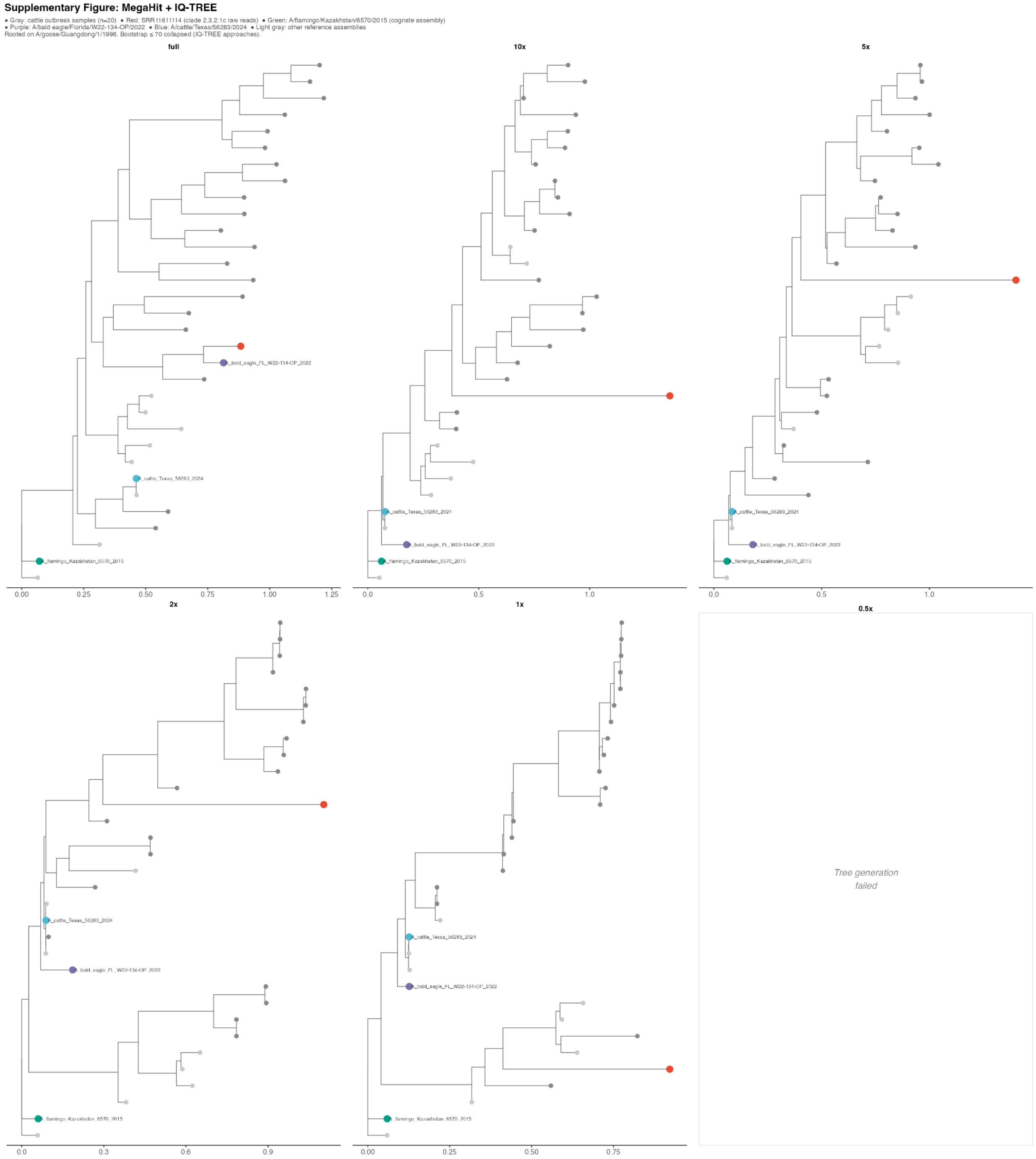


**Supplementary Figure 2. (see above)**

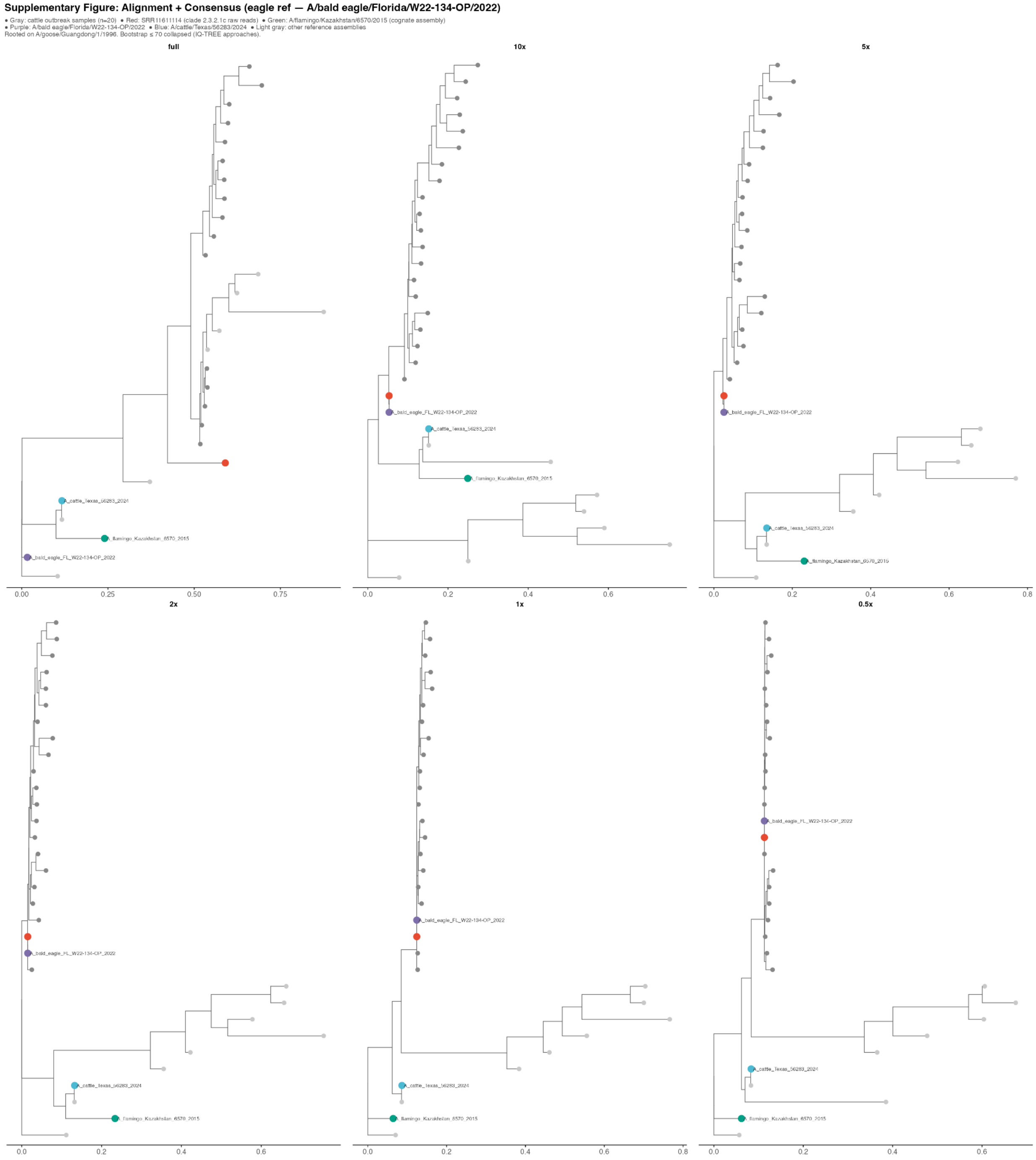


**Supplementary Figure 3. (see above)**

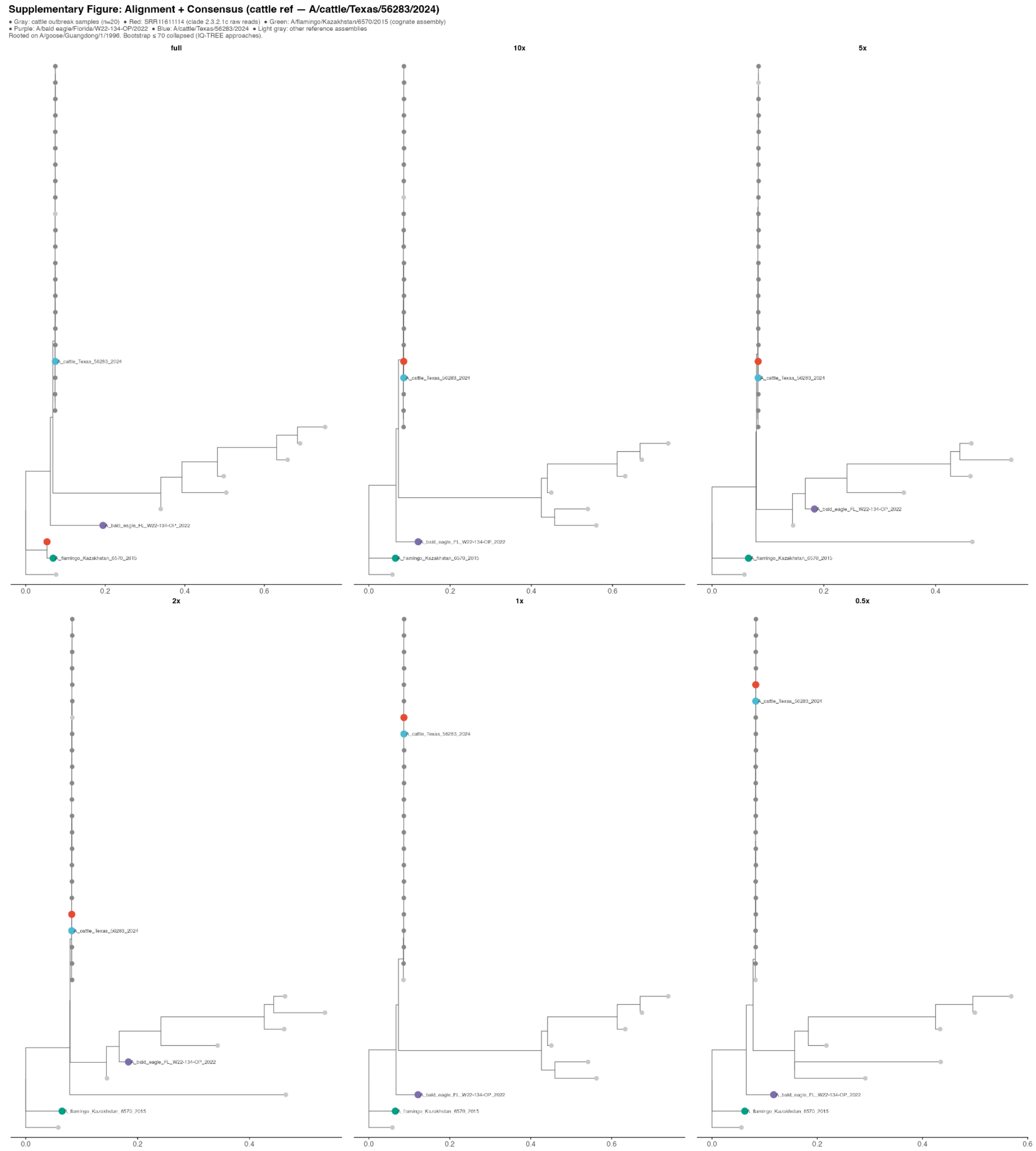


**Supplementary Figure 4. (see above)**

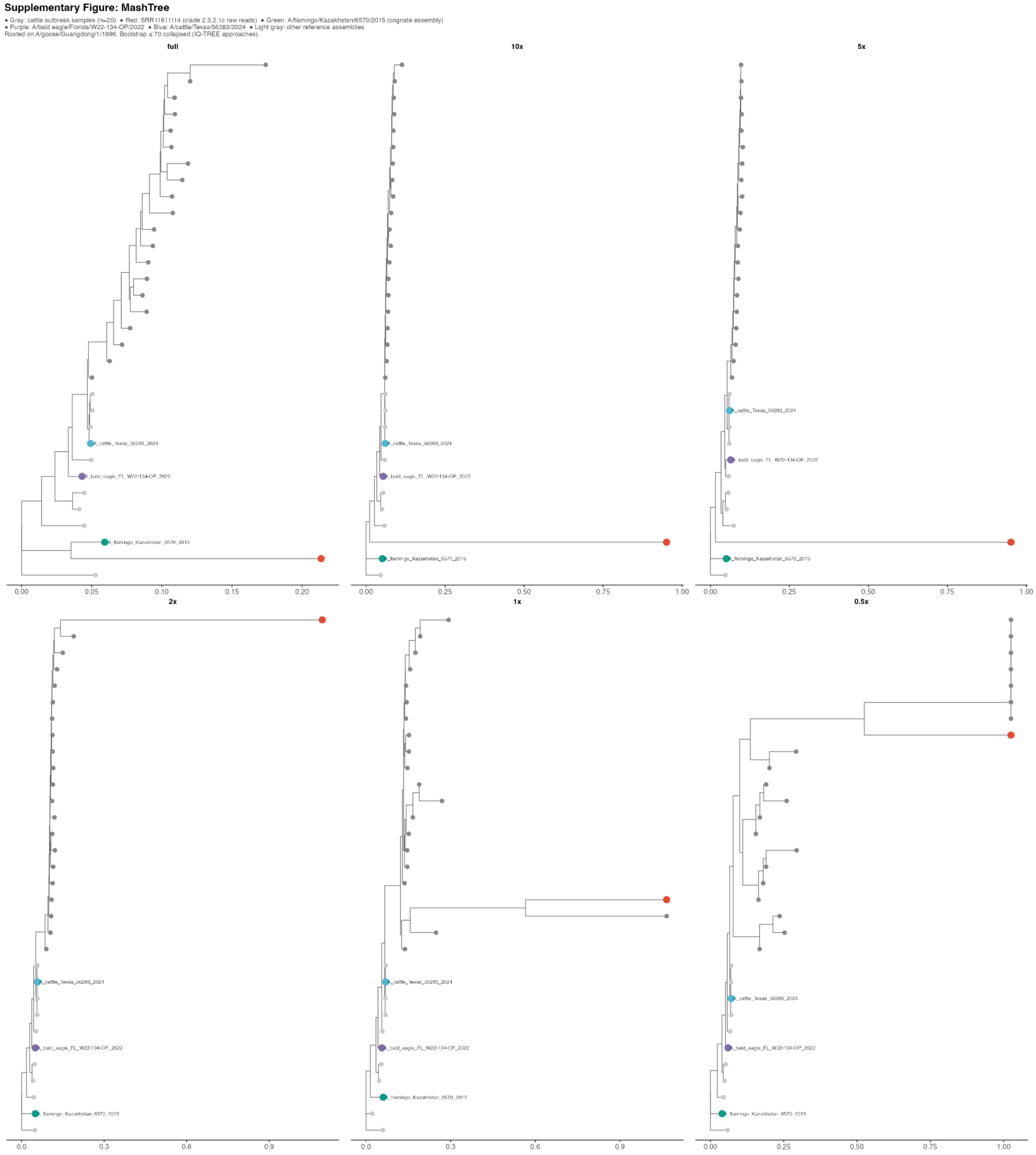


**Supplementary Figure 5. (see above)**

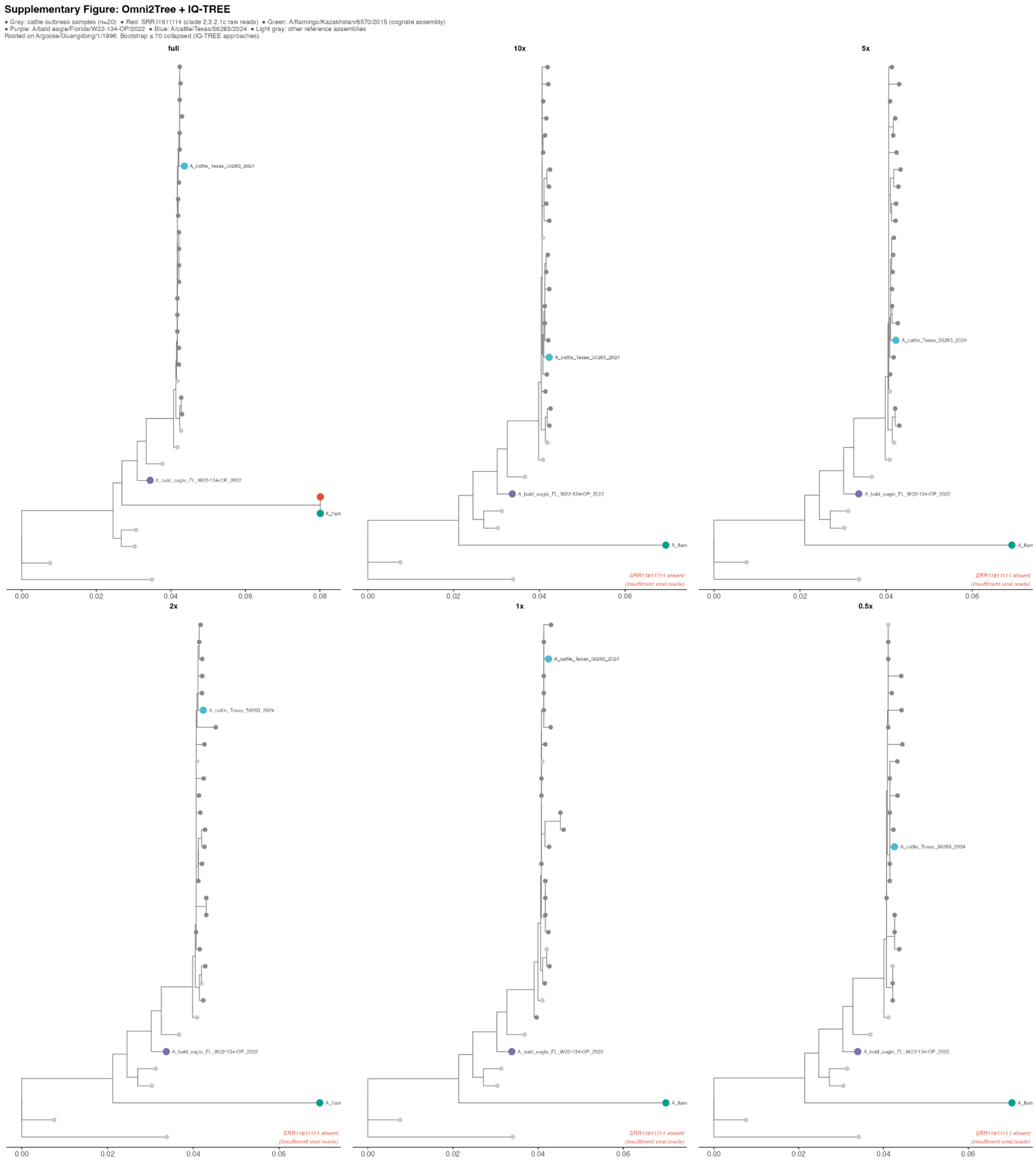


**Supplementary Figure 6. (see above)**

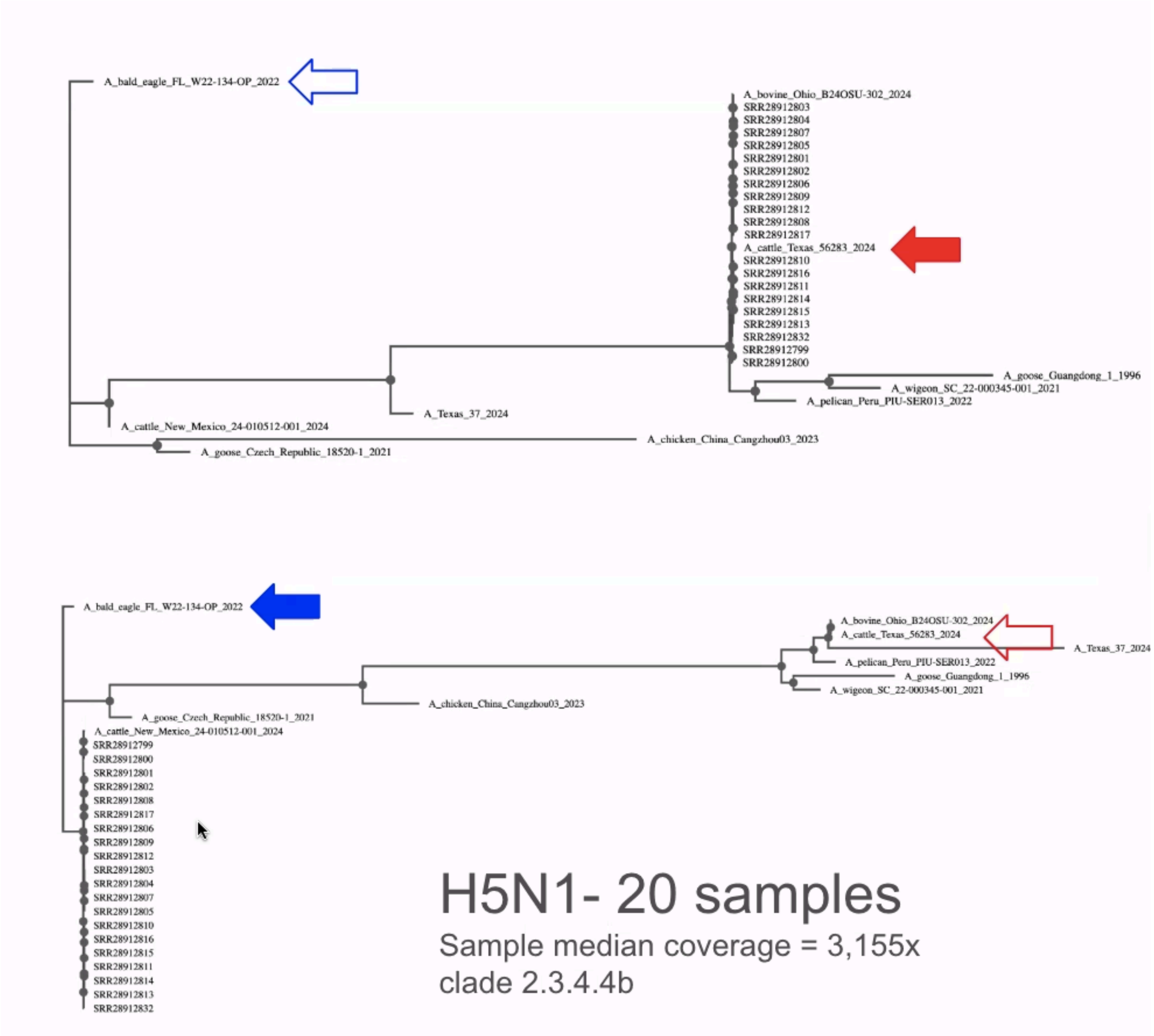


**Supplementary Figure 7. Reference genome choice substantially alters H5N1 phylogeny inferred by alignment + consensus.** Maximum-likelihood trees (IQ-TREE, GTR+G) inferred from reference-guided consensus sequences of 20 outbreak samples and 10 reference strains at full coverage, using two reference genomes for BWA-MEM alignment and bcftools consensus calling: A/cattle/Texas/56283/2024 (top; red arrow) and A/bald eagle/Florida/W22-134-OP/2022 (bottom; blue arrow). Filled arrows indicate the reference genome used for each tree. Sample clustering and branch lengths differ substantially between the two conditions despite identical input reads, demonstrating that consensus-based phylogenetics inherits systematic bias from reference genome choice. Scale bars represent substitutions per site.

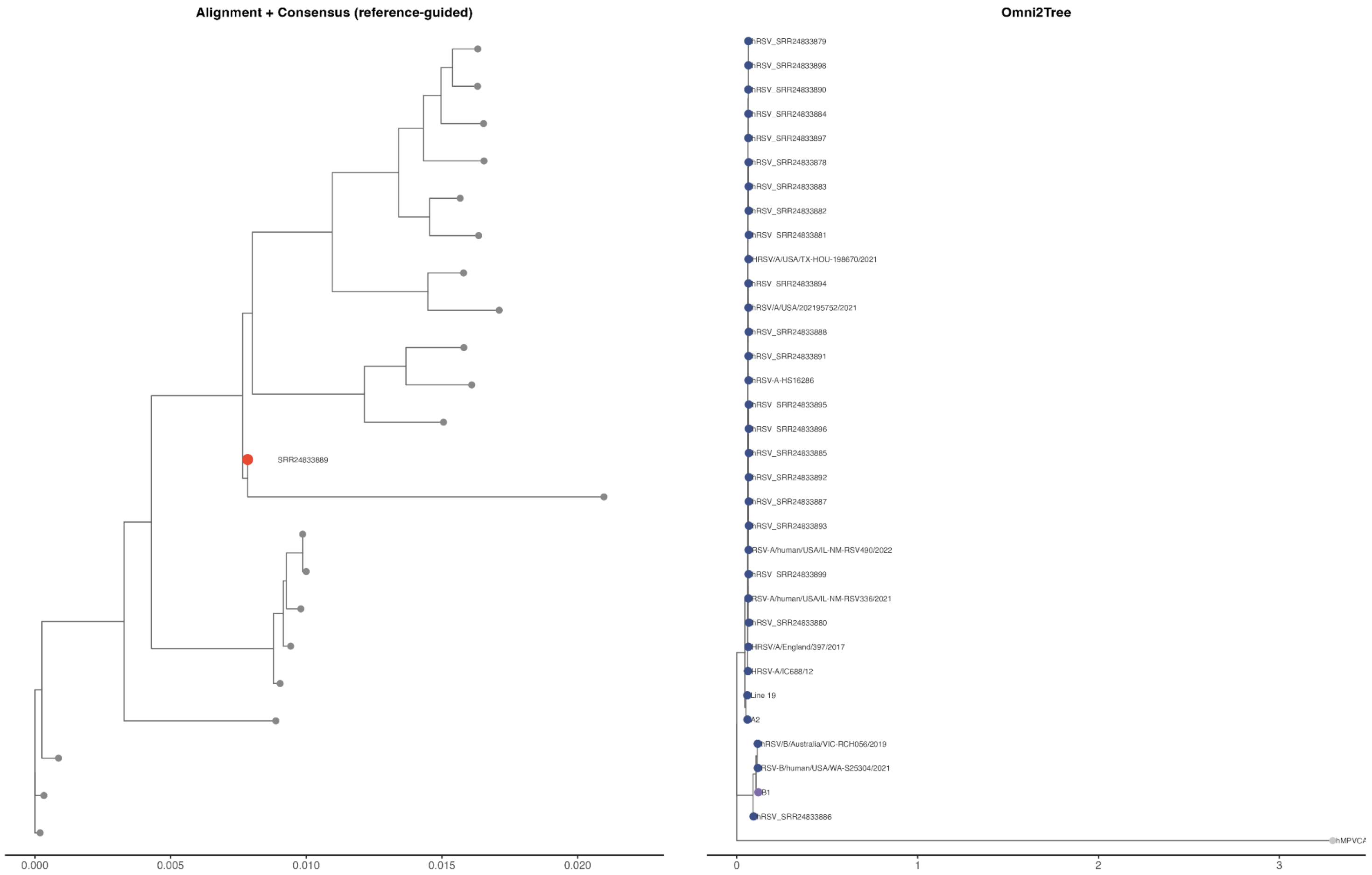


**Supplementary Figure 8. Phylogenetic trees from long-read RSV/A samples (Oxford Nanopore). (A)** Reference-guided consensus tree inferred from 22 RSV/A samples (BioProject PRJNA980575) aligned to an RSV/A reference genome with minimap2, followed by consensus sequence construction with bcftools and tree inference with IQ-TREE (GTR+G, 1,000 ultrafast bootstrap replicates). All 22 samples are placed in the tree regardless of read content, including SRR24833889 (red), which contains zero reads mapping to any RSV reference genome. **(B)** Omni2Tree tree inferred from the same 22 raw ONT read sets using a database containing RSV/A and RSV/B reference strains, with IQ-TREE inference on the concatenated amino acid alignment (LG+G, 1,000 ultrafast bootstrap replicates). SRR24833889 was excluded from the tree by Omni2Tree's quality threshold, as no reads mapped to RSV marker genes. All 21 remaining samples were placed within the RSV/A clade and outside the RSV/B clade, confirming that subgroup assignment is determined by read content rather than database composition. Gray circles = RSV/A samples; red circle = SRR24833889 (zero RSV reads); dark blue circles = RSV/A reference strains (labeled); purple circles = RSV/B reference strains; light gray circle = hMPV outgroup (Omni2Tree only, used for rooting).

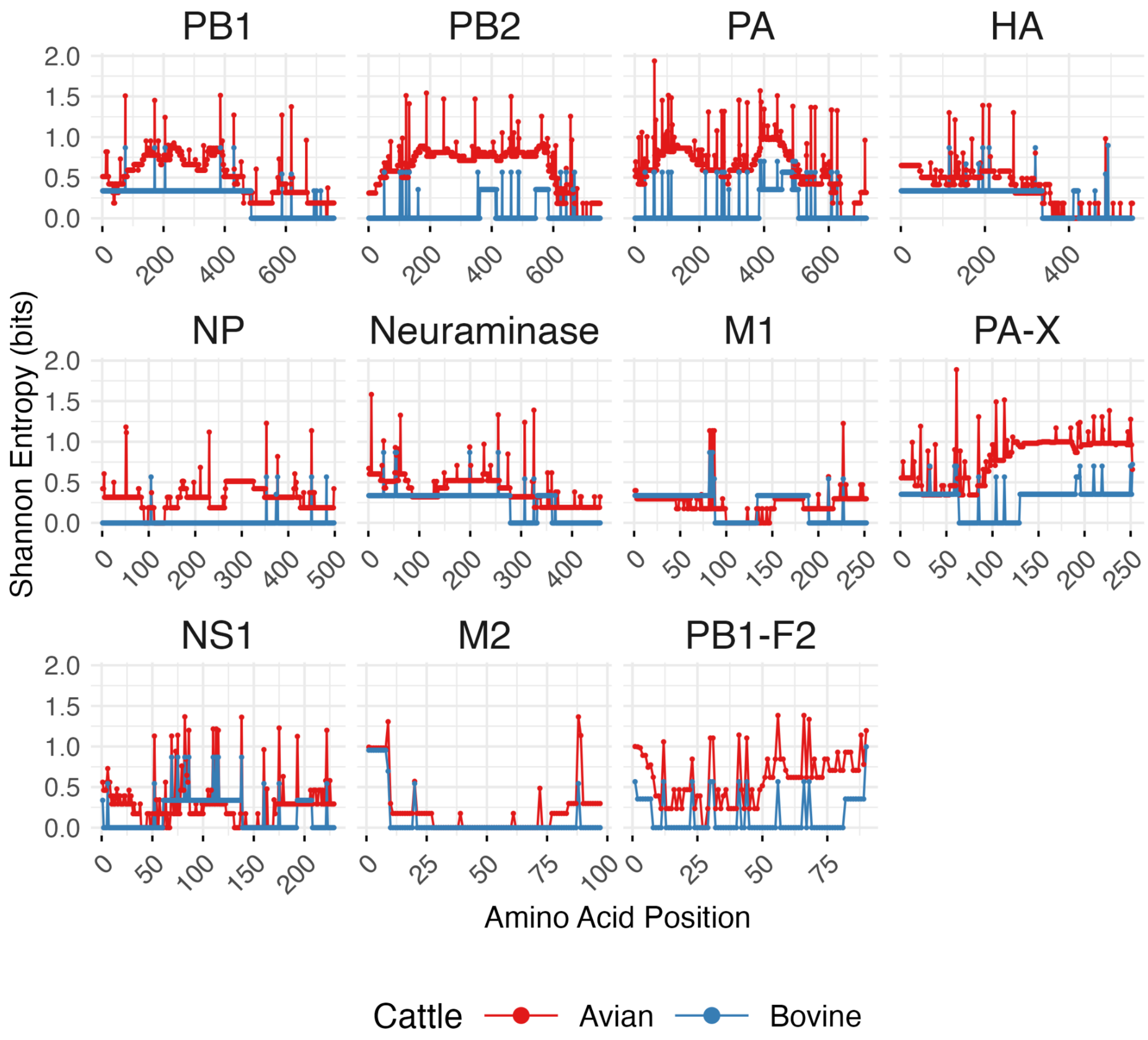


**Supplementary Figure 9. Entropy analysis for all H5N1 genes from the 2024 US cattle outbreak samples.** *Per-position Shannon entropy across all H5N1 proteins, calculated relative to the positions of the A/chicken/Egypt/08102s-NLQP/2009(H5N1) reference strain (Accessions PP395605.1 -PP395612.1). Shannon entropy is expressed in bits; a value of 0 indicates complete conservation, and the theoretical maximum for 20 amino acids is 4.32 bits.*

### Supplementary Note 1

#### Omni2Tree reveals strain-level phylogeny in complex metagenomic communities

Beyond single-pathogen analysis, Omni2Tree extends to metagenomic samples containing mixed microbial populations, a context where assembly is difficult[23,65] and taxonomic classifiers report presence without phylogenetic context. Omni2Tree for metagenomics operates by aligning input sequencing reads from environmental samples to reference gene markers. Omni2Tree then identifies the closest reference strains that are present in the input sample. This is done by selecting those reference strains with sufficient mapped reads (default: 2 reads per gene). For each identified reference, a consensus sequence is generated and combined with all reference sequences to produce a multiple sequence alignment, which is subsequently used to infer a phylogenetic tree (see **Figure 1** and **Methods**).

Omni2Tree metagenomics analysis can be performed as a two-tier approach, depending on the interest or prior knowledge of the user. This starts with a broad placement to characterize unknown communities and is followed by a finer tier round, which is a more focused analysis given a certain strain to delineate. The choice of reference database determines both the taxonomic scope and phylogenetic resolution of the analysis. For samples of unknown composition, we recommend using a phylogenetically diverse reference database.

We benchmarked the metagenomics mode of Omni2Tree on the CAMI Strain Madness challenge datasets[29,66]. This includes simulated complex bacterial communities with multiple co-occurring strains per species. Specifically, we constructed a small diverse database (**Supplementary Table 10**), containing 91 reference taxa strategically sampled from the OMA ortholog database[67] to represent major bacterial lineages across 17 phyla, including Proteobacteria (10 species spanning Alpha-, Beta-, Gamma-, Delta-, and Epsilon-subdivisions), Firmicutes (10 species), Actinobacteria (6 species), Cyanobacteria (7 species), Bacteroidetes (5 species), and representatives from Spirochaetes, Chlamydiae, Chlorobi, Chloroflexi, Deinococcus-Thermus, Aquificae, Acidobacteria, Fusobacteria, Nitrospirae, Planctomycetes, and Verrucomicrobia.

Using this reference database, Omni2Tree placed metagenomic reads from two CAMI datasets into a broad phylogenetic context (**Supplementary Note Figure 1A**). This exploratory analysis detected representatives from 3 bacterial families (*Escherichia*, *Staphylococcus*, and *Streptococcus).* Omni2Tree also reports per-read classification, identifying all the expected six bacterial orders present in the samples (Bacillales, Propionibacteriales, Enterobacterales, Bacteroidales, Eubacteriales, Lactobacillales), even with such a limited reference set. While this analysis reveals which major taxa are present, the limited number of references per genus constrains strain-level resolution. For example, all *E. coli* reads were mapped to two references regardless of how many distinct strains are actually present in the sample. The inferred phylogeny in **Supplementary Note Figure 1A** shows that read populations from the same biological species cluster together across datasets (e.g., D0:ECOSM and D1:ECOSM form a clade, and the same for D0:STAA2 and D1:STAA2), demonstrating phylogenetic consistency independent of sample origin.

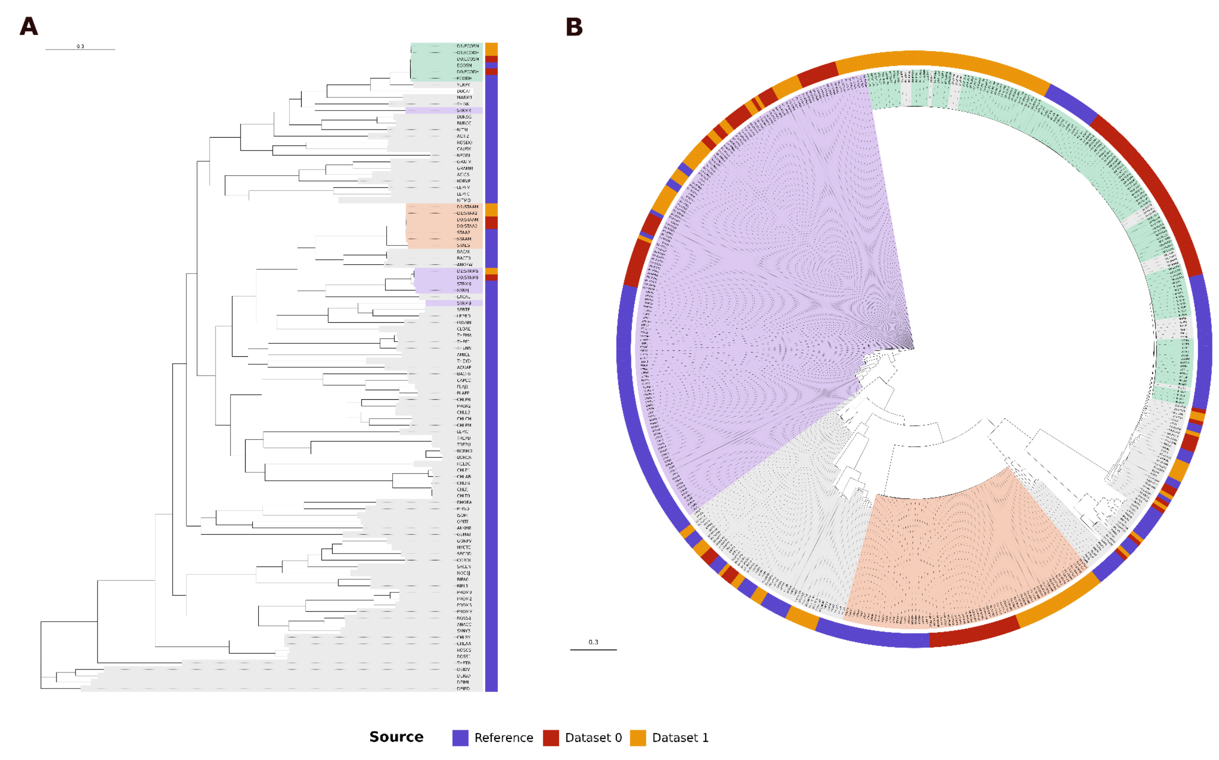


**Supplementary Note Figure 1. a)**: **Omni2Tree exploratory analysis** includes 91 bacterial reference genomes from the OMA ortholog database, and the CAMI2 strain madness datasets 0 and 1. The Maximum-likelihood phylogenetic tree inferred using IQTREE. Reference taxa are indicated by the OMA 5-letter species codes (e.g., ECOSM (Escherichia coli strain SMS-3-5 SECEC), ECODH (ECODH - Escherichia coli strain K12  DH10B), STAA2 (Staphylococcus aureus strain JH1). Metagenomic read populations are labeled as "[dataset]:[reference]", where the prefix (D0 or D1) indicates dataset origin and the suffix (5-letter code) indicates the closest reference species to which reads phylogenetically cluster. Background strips indicate sequences belonging to bacterial species/genera of interest. References without metagenomic matches (e.g., NEORI and NITMO) indicate taxa absent from the analyzed samples. Strain-level resolution is evident where multiple reference strains exist. E. coli populations separate into ECOSM and ECODH clades, while S. aureus populations separate into STAA2 and STAAM lineages. The scale bar represents substitutions per site. **b)** **Omni2Tree focus analysis**, using a dataset of a diverse range of strains. Maximum-likelihood phylogenetic tree generated with Omni2Tree for the CAMI2-specific analysis, containing 193 bacterial strains from the OMA ortholog database, and the CAMI2 strain madness datasets 0 and 1. The outer ring encodes source (Reference, Dataset 0, Dataset 1), while internal tip-background sectors indicate taxa of interest.

Based on the sample placements, users can then perform a targeted analysis restricted to the relevant lineage, enabling higher strain-level resolution within the clade of interest. In the case the study focuses on a specific taxon, such as *E. coli*, this targeted analysis can be performed directly. To demonstrate this targeted approach, we constructed a CAMI-specific reference database using Omni2Tree containing 193 bacterial strains from the OMA ortholog database[67], enriched for taxa expected in the CAMI Strain Madness datasets. Note that if the strain of interest is not in the OMA reference database, an ad hoc set of gene markers could be generated using OMA standalone[68] or FastOMA[69] (Step 1 in **Figure 1**). In this case, the reference gene marker dataset included 41 *Escherichia coli* strains, representing diverse pathotypes (with OMA species code EHEC, ETEC, EPEC, ExPEC, UPEC) and laboratory strains; 23 *Staphylococcus aureus* strains, including MRSA and MSSA lineages; 17 *Streptococcus pneumoniae* serotypes; and multiple strains for *Streptococcus suis* (7 strains), *Enterococcus faecalis* (3 strains), *Shigella* species (7 strains), and other taxa.

Using the 193-strain targeted database, Omni2Tree achieved substantially higher resolution than the exploratory analysis. The resulting phylogeny contained approximately 465 tips: 193 reference strains in addition to 272 read subpopulations partitioned from the two CAMI metagenomic datasets. The phylogenetic tree (**Supplementary Note Figure 1B**) illustrates this strain-level resolution, with reference strains shown in blue and anonymous read subpopulations colored by source dataset (Dataset 0 in red, Dataset 1 in orange).

Each subpopulation represents a distinct cluster of reads assigned to a specific reference strain, for which a consensus sequence is inferred. Omni2Tree detected strain-level populations corresponding to 131 reference strains in Dataset 0 and 141 in Dataset 1, with 127 strains (>90%) detected in both datasets (**Supplementary Note Table 1**). The strain-level resolution arises from the capabilities of Omni2Tree and the granularity of the reference database. Specifically, all 41 *E. coli* strains were matched by distinct read subpopulations, as were all 23 *S. aureus* strains and all 17 *S. pneumoniae* strains. This demonstrates that Omni2Tree can partition reads from closely related strains when appropriate references are available, a resolution that would be impossible with a single reference per species. Read subpopulations consistently cluster with their matched reference strains. When multiple strains of a species are detected, they form species-specific clades reflecting underlying phylogenetic relationships. The *E. coli* clade, for example, shows distinct clustering of EHEC, ETEC, and K-12 lineages, with read subpopulations from both datasets mapping to expected positions within this clade structure. The 18 strains detected in only one dataset likely reflect genuine compositional differences between the CAMI simulations. Four strains were unique to Dataset 0, including three *Streptococcus gallolyticus* strains and one *S. macedonicus* strain. Fourteen strains were unique to Dataset 1, predominantly Lactobacillaceae species (*Lactobacillus*, *Lactococcus*, *Limosilactobacillus*) (**Table 1**). While the CAMI challenge does not provide strain-level ground truth, these differential detections are consistent with expected variation in simulated community composition across datasets.

| **Supplementary Note Table 1:** Species-level strain detection summary (targeted database). Selected species showing strain-level detection across CAMI datasets using the 193-strain targeted database. | | | |
|---|---|---|---|
| **Species** | **Strains in DB** | **Dataset 0** | **Dataset 1** |
| *Escherichia coli* | 41 | 41 | 41 |
| *Staphylococcus aureus* | 23 | 23 | 23 |
| *Streptococcus pneumoniae* | 17 | 17 | 17 |
| *Streptococcus suis* | 7 | 7 | 7 |
| *Streptococcus thermophilus* | 4 | 4 | 4 |
| *Shigella flexneri* | 3 | 3 | 3 |
| *Enterococcus faecalis* | 3 | 3 | 3 |
| *Streptococcus salivarius* | 3 | 3 | 3 |
| *Streptococcus gallolyticus* | 3 | 3 | 0 |
| *Limosilactobacillus reuteri* | 3 | 0 | 3 |
| *Klebsiella pneumoniae* | 2 | 2 | 2 |
| *Enterococcus faecium* | 2 | 2 | 2 |
| *Lactobacillus johnsonii* | 2 | 0 | 2 |
| *Lactococcus garvieae* | 2 | 0 | 2 |
| Total unique strains | 193 | 131 | 141 |
| Common to both datasets | — | 127 | 127 |

To the best of our knowledge, no other tool performs strain-level phylogenetic analysis on metagenomic datasets (such as the CAMI dataset). Thus, we benchmarked Omni2Tree's classification performance against Kraken2, even though Omni2Tree is designed for phylogenetic inference rather than taxonomic classification. Kraken took 2 minutes to perform the classification with a max memory usage of 16.5GB. On the other hand, Omni2Tree took 73 minutes using less than 8.8GB to generate a consensus multiple sequence alignment, followed by tree inference using IQtree, taking 110 minutes using 42 CPUs using 5.20 GB. To evaluate Omni2Tree's performance for metagenomics classification, we calculated the F1 score for all reads. Omni2Tree provides a much higher read-based F1 score compared to Kraken2 when evaluated at the species level (0.63 vs 0.33, the lowest rank provided by the CAMI benchmark) and at the genus level (0.94 vs 0.70) (**Supplementary Table 11**). However, the recall(**Methods**) of Omni2Tree is lower compared to kraken2. This is because kraken2 uses the vast RefSeq genomic dataset of more than 20,000 species, but Omni2Tree in this experiment uses a limited number of species (91) as its reference gene markers. Omni2Tree benefits from the highly accurate read alignment (with minimap) to gene markers, resulting in high precision compared to the k-mer-based approach implemented in Kraken.

We also evaluated how many of the expected species and genera are present in the metagenomic sample. The species-based F1 score of Omni2Tree is much higher than that of Kraken 2 on the two datasets (0.349/0.28 vs 0.0059/0.0053). The same stark difference exists at the genus level (0.509/0.41 vs 0.0069/0.0065) (**Supplementary Table 11**). Kraken results suggest that the sample harbors more than 7,000 species, whereas only 26 species are expected based on the true set. In other words, Kraken suffers from very high false positives in detecting species in the sample (see the ROC curve of Kraken2 for species detection in **Supplementary Figure 9**). On the other hand, Omni2Tree shows a lower recall due to the limited number of reference species. However, this can be addressed by including the set of species that is under study for the experiment, which is straightforward for many clinical applications, such as the study of specific bacteria.

In summary, we showed Omni2Tree's flexibility for metagenomic applications via a two-tier approach, where the exploratory approach determines whether major lineages are present, and the targeted approach with custom databases then enables strain-level resolution, e.g., for tracking S. aureus transmission in a hospital setting. While tools like Kraken2[26] rapidly report species presence, Omni2Tree provides evolutionary context and their phylogenetic relationships. This information is critical for epidemiological applications where strain identity determines pathogenic potential, and for ecological studies where population structure informs community dynamics[70,71]. The ability to process multiple samples simultaneously further enables tracking of strain dynamics across time series, treatment conditions, or geographic locations.

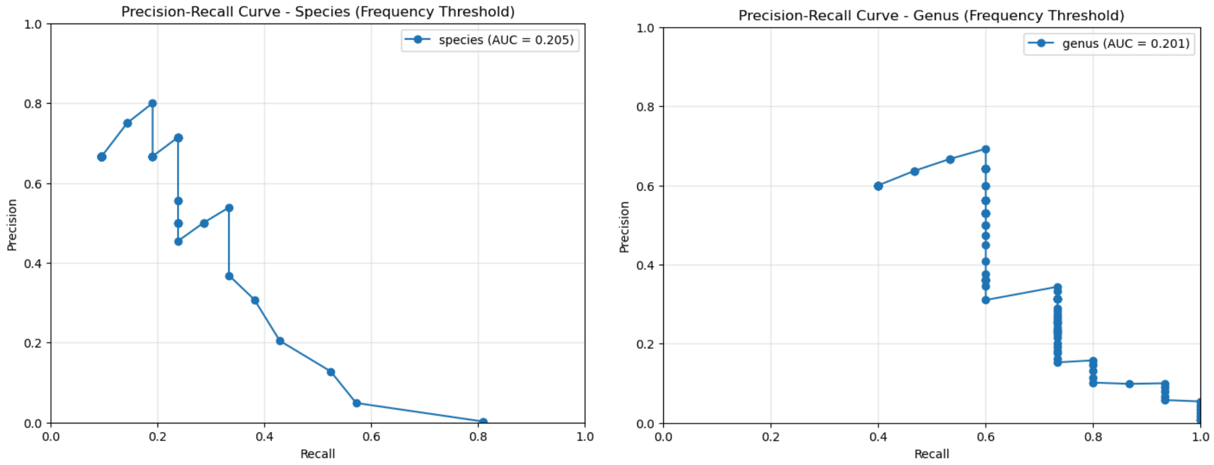


**Supplementary Figure 9**. ROC curve of kraken for species detection.
